# Naturalistic viewing conditions can increase task engagement and aesthetic preference but have only minimal impact on EEG Quality

**DOI:** 10.1101/2021.09.18.460905

**Authors:** Dominik Welke, Edward A. Vessel

## Abstract

Free gaze and moving images are typically avoided in EEG experiments due to the expected generation of artifacts and noise. Yet for a growing number of research questions, loosening these rigorous restrictions would be beneficial. Among these is research on visual aesthetic experiences, which often involve open-ended exploration of highly variable stimuli. Here we systematically compare the effect of conservative vs. more liberal experimental settings on various measures of behavior, brain activity and physiology in an aesthetic rating task. Our primary aim was to assess EEG signal quality. 43 participants either maintained fixation or were allowed to gaze freely, and viewed either static images or dynamic (video) stimuli consisting of dance performances or nature scenes. A passive auditory background task (auditory steady-state response; ASSR) was added as a proxy measure for overall EEG recording quality. We recorded EEG, ECG and eyetracking data, and participants rated their aesthetic preference and state of boredom on each trial. Whereas both behavioral ratings and gaze behavior were affected by task and stimulus manipulations, EEG SNR was barely affected and generally robust across all conditions, despite only minimal preprocessing and no trial rejection. In particular, we show that using video stimuli does not necessarily result in lower EEG quality and can, on the contrary, significantly reduce eye movements while increasing both the participants’ aesthetic response and general task engagement. We see these as encouraging results indicating that – at least in the lab – more liberal experimental conditions can be adopted without significant loss of signal quality.

**Highlights:**

- We assess trade-offs between EEG quality and behavior for common task constraints
- Investigated constraints: fixation task vs. free gaze, and video vs. static stimuli
- Measures: EEG quality, aesthetic preference, task engagement, gaze and heart rate
- Video stimuli reduce eye movements, increase engagement and do not affect EEG SNR
- Fixation task only slightly increases SNR and does not influence ratings

## 1 Introduction

Empirical research often requires a careful trade-off between experimental control and ecological validity. Perhaps nowhere is this balancing act more apparent than when investigating the electrophysiological correlates of higher cognitive functions. Many efforts are taken to design rigorously controlled stimuli that do not introduce potential confounds, and to avoid sources of exogenous and endogenous measurement noise that might obscure the neuronal signals of interest. This has led to the development of a set of experimental constraints that are applied almost canonically in human neurophysiology, particularly in the visual domain. Most prominent amongst these is the nearly ubiquitous fixation task, implemented to reduce the effects of eye movements in electroencephalographic (EEG) signals. Almost as common is the use of static images rather than video or otherwise dynamic visual material. Nevertheless, the last years have seen a movement toward more naturalistic experimentation, with some research areas driving the development of increasingly liberal experimental paradigms. But whereas naturalistic dynamic stimuli have become quite common in the functional magnetic resonance imaging (fMRI) literature (e.g. Hasson, 2004; Hasson & Honey, 2012; Isik & Vessel, 2021; Vodrahalli et al., 2018), the adoption of video stimuli and naturalistic viewing conditions in electroencephalography and magnetoencephalography (MEG) studies is often met with concerns about data quality.

While there can be very good reasons to apply a fixation task or refrain from presenting dynamic visual stimuli, the decision to do so is often a mere cautious default based on common practice, even when it is unclear whether these restrictions might affect the behavior of interest. On the other hand, few researchers who do use naturalistic stimuli, like movies, have systematically compared ‘traditional’ lab experiments with naturalistic experiences to (a) investigate the benefits of naturalistic scenes, for example, in terms of participant engagement, or (b) establish the SNR benefit of static visual scenes over moving images.

Here we seek to inform decisions about experimental design by systematically investigating the effects of relaxing experimental constraints related to naturalistic viewing in an EEG study of visual aesthetic preference. We present participants with either static pictures or dynamic movie stimuli, viewed either under attempted fixation or with free gaze. We then quantify the effects of these two canonical experimental constraints on task relevant behavior, engagement and EEG recording quality.

### 1.1 Relaxing experimental constraints to study naturalistic behaviors

Allowing participants to actively explore stimuli, and adopting the use of time varying visual stimuli in EEG studies would be beneficial for a growing number of research questions related to naturalistic behavior.

In particular, the neuroscientific study of aesthetic experiences would benefit from greater flexibility in experimental design. While fMRI research has identified a set of brain areas and networks involved in aesthetic processing (Chatterjee & Vartanian, 2014; Vessel, 2020), the electrophysiological correlates of aesthetic processing are to date poorly understood. This is unfortunate because the superior temporal precision of EEG and MEG is well suited to investigate the fast neuronal dynamics of aesthetic processing, and the relative mobility of EEG devices could even allow for investigations of aesthetic encounters *in situ*. Yet visual aesthetic experiences in the real world often involve open-ended exploration of highly complex artistic or natural objects. For example, many works of visual art are too large to be captured in one glance and rely on eye, head, or even body movements for active exploration. Other art forms present the observer with moving images or inherently rely on temporal information (e.g. performances, dance, film and time-based media). In addition, the context in which art is encountered matters (see Tinio et al., 2013), and sterile laboratory environments and rigorous experimental constraints might lower the likelihood of strongly moving aesthetic experiences (Brieber et al., 2015; Brieber et al., 2014), thereby negatively impacting statistical power. Investigating the electrophysiological correlates of moving aesthetic experiences would thus benefit from relaxing typical constraints on EEG paradigms.

Beyond empirical aesthetics, such constraints also present problems for other research domains. Static visual stimuli are clearly insufficient for studying other time- or movement-dependent psychological processes, such as motion perception, human movement observation, error detection tasks, and visual narrative. Enforcing fixation is problematic for mental phenomena in which active visual exploration of a stimulus is required, such as visual search, scene perception, the study of reference frames, visual coordination, wayfinding, and decision paradigms that involve foraging. While there are approaches that may allow one to investigate components of these processes within more rigorous frameworks (e.g. using sequential presentation of static pictures), such paradigms remain an approximation and might distort key aspects of the processes of interest. As the fixation task is mostly specific to EEG and MEG studies, it furthermore hinders straightforward comparison with research done using fMRI, fNIRS or purely behavioral methods. Hence, the *a priori* prohibition of the use of moving images and free gaze limits the questions that can be tackled using MEG and EEG, especially in the context of spontaneous or naturalistic behavior.

As stated above, some research areas have already begun developing more liberal experimental paradigms for EEG, though wider adoption of these methods has been limited. In visual neuroscience, the fixation task has been at least partly abandoned when studying scene exploration and visual search (Dias et al., 2013; Kamienkowski et al., 2012; Kaunitz et al., 2014) or reading (Dimigen et al., 2011). Video, moving dot, or other dynamic stimuli are used by researchers interested in motion observation and embodiment (e.g. Heimann et al., 2014), or in hyperscanning studies investigating correlation of neuronal activity between participants that see the same stimuli (Dikker et al., 2017; Dmochowski et al., 2014). There are also a growing number of mobile EEG, MEG, and mobile brain body imaging (MoBI, see Gramann et al., 2014; Makeig et al., 2009) studies in which people’s neuronal responses are recorded during walking (Gwin et al., 2011) and heading changes (Gramann et al., 2021), while navigating complex environments (Debener et al., 2012), or during social interaction (Dikker et al., 2021; Dikker et al., 2017). To address the potential loss of data quality, such studies usually try to counteract the introduction of noise by using better recording hardware, by employing advanced techniques to reconstruct or reject noisy data, or by developing entirely new analysis approaches to target features of the EEG data that are less susceptible to specific sources of noise (e.g. saccade related potentials (Ehinger & Dimigen, 2019; Ossandon et al., 2010) for free viewing, or inter-subject correlation (ISC Ayrolles et al., 2021; Dmochowski et al., 2012) or representational similarity analysis (RSA Kriegeskorte, 2008) for less controlled stimuli). Another approach would be to rely on invasive recordings, such as intracranial EEG, that offer much higher signal quality. However, this is only possible with patients in a hospital and research has shown that even intracranial recordings are susceptible to common sources of noise (Ball et al., 2009; Kovach et al., 2011).

To accelerate the acceptance of new paradigms, we sought to explicitly assess the effect of fixation task and still vs. dynamic video material. Therefore, we systematically investigated electrophysiological signal quality together with a wide variety of behavioral measures targeting potential sources of noise and the participants’ task engagement. We asked participants to view scenes from landscapes and dance performances and to rate each stimulus for both aesthetic appeal and their state of boredom while watching it. The scenes were presented either as dynamic video clips or static pictures, and participants observed them either with unconstrained gaze or under attempted fixation in a fully-crossed, within-subject factorial design. Rather than tracking specific correlates of a given cognitive process, we focused on EEG data quality as a prerequisite to being able to detect such correlates. We expected EEG signal quality to be impacted by the viewing task and by motion modality (still vs. video). At the same time, relaxing these constraints might have a positive effect on behavioral or physiological responses of interest and we sought to quantify the magnitude of these effects and weigh them against each other. Therefore, we also investigated the effects of fixation task and still vs. dynamic video material on aesthetic appeal, task engagement, eye movements and heart rate – measures that are of interest in the study of aesthetic preference and higher cognition in general, but that likely also impact each other in important ways (see below). The content condition (dance or landscape stimuli) functioned both as a control (as the expected rating behavior was well characterized in earlier studies, see below), and to assess the generalizability of any observed effects on EEG data quality.

### 1.2 Quantifying EEG data quality

How can we measure the effects of experimental constraints on EEG data quality? Distortions in EEG recordings can be categorized by their origin into either physiological (endogenous) noise such as eye movements, heart beats, muscle activity (e.g. swallowing, chewing) and task-irrelevant brain activity, or environmental (exogenous) noise such as linenoise, touch/shock on the sensors, electrode impedance and issues with broken sensors or cables. The standard approach is to detect and remove noisy segments or components from the data. EEG preprocessing routines identify broken or noisy EEG channels by employing metrics such as the overall signal amplitude or the correlation with neighboring channels (Luck, 2014), and more specific sources of noise are typically addressed by a researcher’s individual processing pipeline. Visual inspection remains a common way to identify bad data segments that can be very sensitive if done by experts, but this procedure is time-consuming, has low reproducibility, and can introduce (unintended) bias (Rosenthal, 1966). Many advanced techniques have been introduced, including independent components analysis (ICA), canonical correlation analysis, (CCA), artifact subspace reconstruction (ASR), or regression with data from additional modalities (e.g. EOG, EMG, accelerometer), and fully automatic cleaning routines have been proposed to tackle the problem of reproducibility and experimenter bias (e.g. Bigdely-Shamlo et al., 2015; Pedroni et al., 2019).^1^

While many techniques have been developed to avoid noise and to clean recorded data, there is a striking shortage in the literature when it comes to quantifying the amount of noise in a EEG data. This might be partly due to the fact that what is considered “noise” at least partly depends on the “signal” one is interested in. The few metrics that have been proposed usually aim to assess the SNR of expected and well characterized evoked responses such as ERPs (Luck, 2014; Luck et al., 2021; Picton, 2011; Wong & Bickford, 1980).

In the present study, however, we needed a more versatile quantitative measure of EEG recording quality that can change over time, thereby allowing for a *within-subject* comparison of different trials or segments of data in a continuous EEG recording, and that is not interfering with the neuronal processes of primary interest in the study. The desired metric has to be sensitive to the many different sources of endogenous and exogenous noise in EEG recordings described above. Such quantitative measures of EEG quality would arguably be a very practical tool for many researchers, even beyond those who have to abandon some level of experimental control in their data collections. Previous methods-focused studies in the MoBI field have employed known and robustly evoked brain responses to systematically compare signal quality across different conditions. For example, Debener et al. used the P300 ERP-response to an auditory oddball task as a proxy measure to quantify EEG quality of a customized, low-cost, mobile EEG system as participants walked around a campus or sat still in an office space (Debener et al., 2012). Unfortunately, however, datasets like these are often collected solely for the purpose of piloting experiments (and are hence under-powered) or during development and testing of recording setups, and are hardly ever published.

Here we utilize the auditory steady-state response (ASSR) as a proxy measure for overall EEG recording quality. We deliberately chose an auditory process to minimize interference with the primary visual task. ASSR is an automatic neuronal response to a periodically modulated auditory stream (Stapells et al., 1984). The ASSR is generated throughout the auditory system with contribution from both brainstem and cortical regions and can be elicited by a broad range of auditory stimuli such as click trains, beats, tone bursts, amplitude modulated sine waves or noise, and frequency modulated signals (Picton et al., 2003). The brain tracks the auditory signal and its response can be quantified from the frequency domain EEG signal as peaks at the stimulation frequency and its corresponding harmonics (Meigen & Bach, 1999; Norcia et al., 2015; Picton et al., 2003). ASSR is stable over time (Van Eeckhoutte et al., 2018) and has been shown to not interfere with visual processing (Keitel et al., 2013) In this study an ASSR-eliciting auditory stream accompanied all trials as a passive background manipulation. The stimulus was optimized to be as unobtrusive as possible, with a relatively low volume adapted to each participant’s individual hearing threshold. It is important to note that there is a difference between the brain response (ASSR *sensu strictu*) and the ASSR signal that is measured at the scalp. If the EEG recording quality drops while an auditory ASSR stream is presented to a listener (i.e. if the ASSR is overlayed by endogenous or exogenous noise), we expect the strength of the continuously measured ASSR signal to decrease. In order to exploit this for our metric, the EEG data collected in this study were only minimally preprocessed and were not cleaned to remove noisy segments.

### 1.3 Measuring aesthetic appeal, engagement and physiological responses under varying task conditions

#### Aesthetic appeal

Our primary behavioral measure of interest is rated aesthetic appeal. It is important to characterize the potential effects of viewing restrictions on aesthetic valuation as previous research has shown that the presentation context matters (e.g. viewing artworks in a museum vs. on a computer screen Brieber et al., 2014). This effect may be partly due to observers’ ability to freely explore the stimuli, as gaze patterns and viewing time have been linked to aesthetic valuation (Mitrovic et al., 2020). To our knowledge, we are the first to investigate potential effects of dynamic video material compared to static stimuli. We also expect higher aesthetic ratings for nature stimuli compared to dance videos (Isik & Vessel, 2019). While aesthetic preferences for specific stimuli can be highly variable across observers (Vessel et al., 2018; Vessel et al., 2012), many studies have shown an overall effect of the content domain on aesthetic ratings, with higher average ratings and greater across-observer agreement for natural landscapes and faces compared to other categories (e.g. Vessel et al., 2018; Vessel & Rubin, 2010). We operationalize our measure as *aesthetic* appeal, but very similar ratings are also collected in other research on preference and reward (e.g. Lopez-Persem et al., 2020). Hence we believe that the results will also apply to more general measures of preference, liking or sensory pleasure.

#### Boredom and task engagement

We sample the participants’ state of boredom in order to operationalize task engagement vs. disengagement. Both disengagement and states of boredom have been consistently related to task-unrelated thought and fluctuations of attention (Raffaelli et al., 2018; Smallwood et al., 2004). We expect lower boredom ratings in the free-viewing task compared to the fixation task, and a moderate negative correlation between boredom and aesthetic preference ratings. To our knowledge, no study to date has collected both aesthetic and boredom ratings at the same time and hence the interplay of these two behavioral ratings is of particular interest. Periods of spontaneous task-unrelated thought, e.g. mind wandering, have been shown to involve the default-mode network (DMN) (Fox et al., 2015), and the few available fMRI studies on boredom also consistently reported activation of this network (Raffaelli et al., 2018). Interestingly, the DMN has also been implicated in aesthetic processing (Vessel et al., 2019; Vessel et al., 2012). Low engagement can also impact behavioral data quality (e.g. false responses or reaction times McVay & Kane, 2012) and EEG recordings. Decreased task engagement marked by periods of mind wandering was shown to negatively affect the strength of ERP components (Kam et al., 2011; Smallwood et al., 2008) and is marked by time-frequency components in the alpha and theta band (Braboszcz & Delorme, 2011; Kam et al., 2021). Further, disengagement with the task at hand might also lead to increased physical restlessness or higher eye blink frequency (thereby introducing artifacts) and increased fatigue.

#### Eye tracking

We recorded eye tracking data both to test whether participants maintained fixation during the fixation task and also to explore whether any of the other experimental conditions significantly impacted the observers’ eye movements. In EEG research, eye movements are mainly regarded as problematic artifacts (Iwasaki et al., 2005) that should be avoided or suppressed. Eyeblinks and larger saccades can be identified in the time domain EEG and would most likely lead to a rejection or reconstruction of data by any of the cleaning routines described above. Smaller fixational eye movements, also called microsaccades, cannot be easily identified from the EEG but might be problematic as well, since they have been shown to induce transient gamma-band responses in the EEG (Yuval-Greenberg et al., 2008) that could be misinterpreted as brain activity. While a fixation task significantly reduces participants’ saccade counts (Otero-Millan et al., 2008), this mainly inhibits voluntary saccades and blinks – microsaccades, and their potential negative effects on EEG signal, also occur under attempted fixation (Otero-Millan et al., 2008; Thielen et al., 2019).

However, eye movements must not be conceptualized as “noise” alone. In fact, they are meaningful goal-directed behavior that controls the visual input stream to the observers brain. Furthermore, previous research indicates that eye movement patterns may carry relevant information in the context of boredom, engagement, and aesthetic processing: viewing times and fixation heat maps are frequently used in behavioral work on empirical aesthetics (e.g. Mitrovic et al., 2020), microsaccades can carry information about internal states during music listening (Fink et al., 2019), and average fixation duration has even been proposed as a general marker for engagement and external focus during visual tasks (Ramos Gameiro et al., 2017).

Previous work from the eye-tracking literature suggests that there might be differences between still and video stimuli in terms of observers’ eye movements — a phenomenon called center bias. This tendency to keep the gaze focused at the center of a visual stimulus during the time of presentation appears to be stronger in videos than in static stimuli (for a review see T. J. Smith, 2013).

#### Heart rate variability

Heart rate (HR) was included in our study on an exploratory basis. Changes in HR have been related to boredom and engagement (Raffaelli et al., 2018) and are in general often investigated when interested in contributions of the autonomous nervous system (ANS), especially in the context of emotional processing (Palomba et al., 1997; Patrick et al., 1993; Vrana et al., 1988; Winton et al., 1984). Eye movements have also been reported to correlate with change of HR and other ANS responses (Liu et al., 2020). Boredom can cause increased heart rate, decreased skin conductance and increased cortisol level, hinting at high arousal and difficulties in sustaining attention (Merrifield & Danckert, 2014). Slowing of the heart beat, on the other hand, has been associated with higher engagement (i.e. increased focus and concentration) (Coles, 1972). Heart rate might thus be a promising measure in both directions of a hypothesized boredom-engagement continuum. Heartbeats are also relevant in the context of EEG quality: heartbeat evoked potentials (HEP) (Schandry et al., 1986) are not typically controlled for in EEG cleaning routines (if not captured by ICA), and hence contribute to the measured and analyzed signal. Importantly research suggests that the amplitude of the HEP can be affected by factors such as attention, arousal, or internal focus (for a review see Coll et al., 2021), which might lead to confounds if these are systematically linked to the experimental conditions.

## 2 Methods

The experimental design and our hypotheses were preregistered (see https://osf.io/bkep4). Any detail of the final methods that differed from the preregistered plan is clearly described and discussed as such. Information regarding participants and recording devices are reported following the standards proposed for M/EEG studies by the Committee on Best Practice in Data Analysis and Sharing (COBIDAS) of the Organization for Human Brain Mapping (OHBM) (C. Pernet et al., 2020; C. R. Pernet et al., 2018). Data were collected from Oct. 2019 through Jan. 2020.

### 2.1 Participants

#### Sample size estimation

Target sample size was estimated via *a priori* power analysis using G*Power software (Faul et al., 2007, version 3.1). Calculation for a fixed effects ANOVA with 8 groups, and a numerator df = 1 (all main effects and possible interactions individually) for an effect size of Cohen’s *f* = 0.5, alpha = .05, and power = .8 resulted in a target sample size of 34 participants. Due to expected dropouts we planned to collect data from at least 40 participants. During the preparation of the manuscript we realized that we had erroneously computed the statistical power for an across-subject rather than a within-subject design as fit for the collected data (we wrongly chose “ANOVA: Fixed effects, special, main effects and interactions”). This lead to a significant overestimation of the required sample size: with the correct design our final sample offers sensitivity to detect effect sizes of Cohen’s *f* > 0.21 with a critical *F* = 2.04 (“ANOVA: Repeated measures within factors” with alpha = .05, power = .8, total sample size = 43, 1 group, 8 measurements, and conservative expectation of 0 correlation among repeated measures).

#### Recruitment

Participants were convenience-sampled from the general public in the Rhein-Main metropolitan region in Germany. Recruitment was performed via advertisement on the institute website and direct mailings to members of an institute hosted participant database (>2000 members, open to everybody to subscribe). Slots were assigned on a first come, first serve basis. Inclusion criteria for recruitment were age between 18-55 years, no impaired hearing, normal or corrected to normal vision and no need to wear glasses during the study (as this might decrease the quality of the eye tracking).

#### Final sample

47 participants enrolled for the experiment and 45 participants finished data collection. Data from 43 participants were included in the final analysis.

Two types of non-systematic and non-reproducible software errors appeared in some of the recording sessions: 1) a sound-driver related error leading to absence of the auditory stimulus, and 2) a screen freeze, probably linked to the interaction of the eye tracking system with the presentation PC. Two recording sessions were aborted when these software errors appeared for the first time, and data were excluded from analysis because a reconstruction of the time of occurrence was impossible. Two further participants were excluded from EEG analysis due to erroneous amplifier settings that resulted in a sampling frequency of only 250 Hz (incongruent with the preregistered design).

Five of the 43 remaining participants had missing trials due to either of the above mentioned software issues but were included in the analysis. From these 43, one participant reported minor neurological problems (peripheral nerve damage after an accident). Nine other participants reported current or past episodes of mental or psychological disorders. One of these nine indicated current and long-term medical treatment.

The participants were between 19 - 52 years old (mean = 27.1 years, std = 7.1 years), 26 of them female, 17 male, and 0 indicated “other.” Participants had received between 9 - 25 years of education (mean = 17.8 years, std = 3.6 years), with the sample showing a strong bias towards highly educated people: 40 out of 43 held the German Abitur or a university degree as their highest qualification. 39 of the participants were right handed, 2 left handed, and 2 ambidextrous (based on self report). Eye dominance was assessed by the experimenter (see below): 33 participants exhibited right eye, 6 left eye, and 4 no eye dominance. Although it was not a criterion for exclusion, we also assessed the participants’ caffeine intake on the day of the experiment: it ranged between 0 - 4.28 mg/kg body weight (mean = 0.84 mg/kg, std = 1.04 mg/kg). 15 out of 43 participants indicated zero caffeine intake before participation. See Supp. Tabs. 2 and 3 for full details on participant demographics.

All participants received monetary compensation of 14 euro per hour and gave their informed written consent prior to participation. The study adhered to the ethical standards of the Declaration of Helsinki and was approved by the local ethics committee (Ethics Council of the Max Planck Society).

### 2.2 Experimental design

This study was laid out as a 2×2×2 fully crossed factorial design with each of the three stimulus factors having two levels: factor 1 – fixation task (attempted fixation or free gaze); factor 2 – stimulus dynamics (dynamic video stimuli or static picture stimuli); and factor 3 – stimulus content domain (dance or landscape scenes). An overview of the design is shown in Fig. 1.

**Figure 1:**
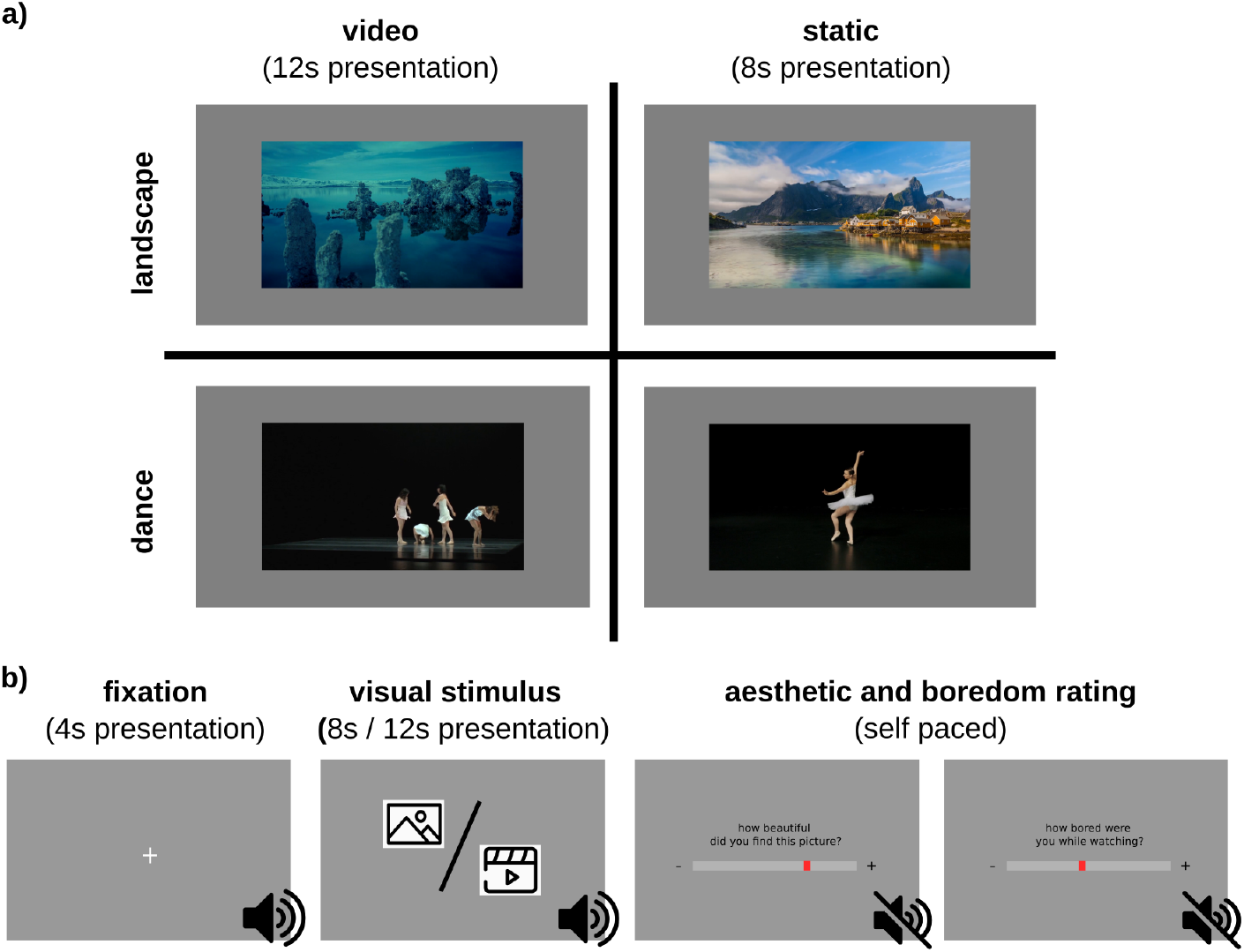
Overview of the experimental paradigm. a) fully crossed 2×2×2 factorial design with either dance or landscape stimuli, and either dynamic video or static picture stimuli; the third condition, either unconstrained gaze or attempted fixation on a fixation dot, is not shown in the Figure. b) general sequence of each of the 80 experimental trials: 4s of fixation dot on grey background are followed by the visual stimulus (8s or 12s respectively). An auditory stream that elicits an auditory steady-state response (ASSR) accompanies these parts. Afterwards, behavioral ratings of aesthetic appeal and state of boredom during the trial are collected (self-paced).

We presented 80 trials per participant in a fully balanced fashion resulting in 40 trials within each level of a given stimulus factor (e.g. video stimuli, regardless of content and fixation task) and 10 trials for each possible combination of factors (e.g. static landscape pictures with fixation task). Rating responses were collected following each stimulus presentation trial.

Participants finished all parts of the study in a single session. Electroencephalographic (EEG), electrocardiographic (ECG), and electromyographic (EMG; data not analysed) data were recorded continuously, together with a marker channel for subsequent epoching. Eye tracking data were recorded in epochs time-locked to each trial start.

#### Task procedure

First the participant answered a set of ancillary questionnaires (see below) in digital form on a laptop computer, while the experimenter prepared the EEG cap. Ocular dominance was determined by the experimenter using a variation of the Porta test (as described in Roth, 2002).

After that the participant entered the cabin, the EEG cap was mounted, and peripheral electrodes (ECG and EMG) were attached. EEG electrodes were filled with electrolyte gel and electrodes were adjusted until impedance were below or close to 10 kOhm. After finishing preparations, the effects of chewing, blinking, and contracting neck muscles on the EEG signal were demonstrated to the participant to raise awareness for these sources of artifacts, along with a demonstration of individual alpha oscillations during a short eyes closed period.

To begin the experiment, the participant’s personalized hearing threshold was determined with a self paced staircase 1-up-1-down task. A short burst of the same auditory stimulus presented in the experiment was used for the threshold detection, starting with an unnoticeable low volume. Participants were instructed to increase the volume until they heard the stimulus for the first time, then decrease volume until they did not hear the stimulus anymore, then change direction again and so forth. After 7 reversals the task ended, and the individual sensory threshold was set as the average of the last 3 reversal intensities.

After the end of the threshold task the experimenter entered the cabin, adjusted the participants’ sitting position and the head rest, and set up the eye tracker. From then on, the participant was asked to keep her position in the head rest and restrain from chewing, swallowing or moving the head during trials.

Instructions for the main experiment were displayed, followed by a training session consisting of a calibration routine for the eye tracker (horizontal and vertical calibration at 5 positions) and four practice trials. The four practice trials contained stimuli from all possible conditions: dance and landscape, video and static, fixation task and free gaze. These stimuli were not presented again in the main experiment. The main experiment then started with a calibration of the eye tracker followed by the first block of trials.

The general sequence of each trial is shown in Fig. 1b. A trial began with 3 s of a blank screen, accompanied by auditory stimulation. Auditory stimulation continued during presentation of a fixation cross (4 s) and presentation of either a video clip (12 s) or static picture stimulus (8 s). Auditory stimulation stopped with the end of the image or video stimuli, and was followed by a blank screen (4 s), a self-paced aesthetic rating (see below), another blank screen (1.5 s), a self-paced boredom rating (see below), and a final blank screen (2 s). A mid-gray background (*rgb*[128, 128, 128]) was used for the entire experiment.

The presentation durations were chosen as a compromise: for video clips, shorter than 12s is quite short for an experience to evolve, while long periods of static pictures might be experienced as boring. EEG analyses were performed on a common time window of 0-8 s (last 4 s of video presentation omitted).

Stimuli were presented in 16 blocks of 5 trials, consistent in modality (dance or landscape, video or static) and observation task (fixation or free gaze). The presentation order of stimuli within the blocks was randomized. The same random order was applied for all participants, yet counterbalanced by reversing it for every other participant. Source stimuli were not repeated until each was shown once (i.e. the first 40 trials consisted of 20 pictures and 20 videos from the 40 different source clips; see below).

At the beginning of each block the type of observation task (fixation or free view) and a counter of the remaining blocks was displayed to the participant. Block conditions were intermixed; the block order was counterbalanced across participants. The eye tracker was re-calibrated at the beginning of every other block. A longer self-paced break was offered after half of the blocks (40 trials).

#### Behavioural measures and questionnaires

First, participants answered a set of questionnaires: basic demographic information (including age, gender, education, and information on mood or neurological disorders), caffeine intake of the day (Bühler et al., 2014), the big-five personality inventory (BFI-2-XS: Rammstedt et al., 2020; Soto & John, 2017), the short boredom proneness scale (sBPS: Struk et al., 2017), the positive negative affect schedule (PANAS-SF: Thompson, 2007), and the Snaith-Hamilton pleasure scale (SHAPS: Snaith et al., 1995) in their German versions.

Ratings of aesthetic appeal and state boredom were collected using a continuous scale with a slider controlled by moving the mouse. A response was logged by clicking the mouse button, but participants could not log a response without first moving the mouse in order to prevent lack of responding. Both the ratings and response times were logged. The ends of the scales were marked with “+” and and each scale was accompanied by the corresponding question: “Wie sehr hat diese Szene Sie angesprochen?” (“How much did this scene appeal to you?”) for the aesthetic rating, or “Haben Sie sich beim betrachten der Szene gelangweilt?” (“Were you bored while watching the scene?”) for the boredom rating respectively. The order of the ratings was fixed with the aesthetic rating always coming first.

In the more detailed task description preceding the study, the aesthetic rating was related to the concept of “being moved” by an aesthetic stimulus (Menninghaus et al., 2015): This psychological construct has been used in several behavioral and MRI studies both by our lab and others (Armstrong & Detweiler-Bedell, 2008; Menninghaus et al., 2015; Silvia, 2009; Vessel et al., 2012). Previous research on boredom typically applies induction paradigms via specific boring tasks or video material (Raffaelli et al., 2018). Since here we are interested in dynamic changes of state boredom over the course of the experiment we apply repeated sampling with a self-report rating analogous to the assessment of aesthetic appeal. In the task descriptions we emphasized that this question was targeting the participant’s state of perceived boredom, not their evaluation of the stimulus or any of its features.

### 2.3 Stimuli

#### Video stimuli

The video stimuli were generated from a larger set of 60 video clips (30 dance performances, 30 landscapes) of 30 s length compiled for a previous study (Isik & Vessel, 2019). These clips were screened by a lab assistant naïve to the purpose of the study: all shots (continuous segments without cuts) of 12 s or longer were identified and 12 s excerpts starting with the first frame after a cut were extracted using Adobe Premiere (Adobe Inc.). If any 30 s clip had no cuts at all or contained shots much longer than 12 s the assistant was instructed to extract 12 s excerpts that did not start in the middle of a fast pan (camera shift) or dance move. This procedure yielded a set of 86 excerpt clips from 47 of the 60 original 30 s clips (0-3 per original clip; 53 dance performances, 33 landscape). From these 86 clips 40 were chosen as stimuli for the experiment (20 landscapes, 20 dance scenes; random choice, with only one clip per source video). 1 of the 40 videos was in grayscale and the remaining 39 were color videos. All video clips had an aspect ratio of 16:9, an initial resolution of 1280×720 px, and were compressed in the same video compression method (H.264).

#### Picture stimuli

The static picture stimuli were still-frames from each of the 40 chosen video stimuli (one picture stimulus per video stimulus). Frames were taken from the last 2 seconds of each clip and had to be free of motion smearing (especially in dance performances). Other than these restrictions, frame selection was based on experimenter choice.

Videos and pictures were scaled not to exceed 20° *deg* of visual angle in the vertical or horizontal dimension, resulting in a stimulus size of 502 x 282 px on the screen.

#### Auditory stream to elicit ASSR

During each trial, pink noise with a continuous 40 Hz amplitude modulation was played to participants via binaural in-ear headphones with a loudness of 35dB SL (i.e. 35 db louder than their individual sensory detection threshold for the stimulus; see above for description of the threshold detection task). The final loudness was controlled via the presentation software. We chose 40 Hz stimulation frequency because ASSR has been shown to be particularly strong in response to this frequency range (Galambos et al., 1981), possibly due to superposition of brainstem and middle latency responses (Bohórquez & Özdamar, 2008).

To generate the auditory stimulus a 30 s pink noise waveform was created using MATLABs *pinknoise* function and dotwise multiplied with an equally long sine wave as the modulating factor; the depth of the amplitude modulation was hence 100%. The stimulus was saved as an uncompressed wav file with 44.1 kHz sampling rate. It can be retrieved from the associated online repository.

Presentation of the auditory stream started 3 s before the first frame of the pre-stimulus fixation cross and ended with the last frame of the visual stimulus, resulting in a total of 15 s auditory stimulation in static picture trials and 19 s in video trials, respectively. In this design the initial transient response period of the ASSR lies outside the analyzed time window of visual stimulation (see below).

### 2.4 Data acquisition and devices

#### Study environment

EEG preparation and main experimental routine took place in an acoustically shielded cabin (model: IAC 120a, IAC GmbH, Germany; internal dimension: 2,74 x 2,54 x 2,3 m). Participants were seated in a chair and placed their chin on a chin rest with forehead support (SR Research Head Support, SR Research Ltd., Canada). The distance between the chin rest and the screen was 72 cm. The height of the desk was adjustable such that participants could sit in a comfortable upright position that they were able to sustain for the time of the study.

The study was run by three different experimenters, with the vast majority (44 out of 47 participants) run by one. The participant could contact the experimenter any time via a room microphone installed in the cabin.

The experiment was run on a PC running 64-bit Microsoft Windows 7.1.7601 service pack 1 (Microsoft Corporation, USA), using PsychoPy3 standalone software (Peirce, 2007, version 3.0.7). Visual stimuli were presented on a 24 inch BenQ XL2420Z screen (BenQ Corporation, Taiwan) with nominal framerate of 144 Hz, resolution 1920 x 1080 px (mirrored for the experimenter), and auditory stimuli were presented using an RME Fireface UCX Audiointerface (Audio AG, Germany) with ER3C Tubal Insert Earphones (Etymotic Research Inc., USA).

#### EEG and peripheral physiology data acquisition

EEG data were collected using a 64 channel actiCAP system with active Ag/AgCl electrodes with no active shielding (Brain Products GmbH, Germany), placed according to extended international 10-20 localization system (Committee, 1958; Oostenveld & Praamstra, 2001) with FCz recording reference and AFz ground. The cap was positioned by centering the Cz electrode on the axes *Nasion* to *Inion* and left ear to right ear. The channel layout included 2 bipolar auxiliary channels for ECG (electrodes placed on the right mid clavicle and lower left rib cage, in correspondence with the II Einthoven’s derivation) and EMG (electrodes placed on left forearm, on top of *Musculus brachioradialis* and *Musculus extensor carpi radialis longus* with a distance of approximately 5-10 cm depending on the participant’s size; EMG data were not analyzed). The skin at electrode sites was cleaned with alcohol in advance.

Signals were amplified using a BrainVision actiCHamp 128 Amplifier with 5 BIP2Aux Adapters with 24-bit digitization and an analogue bandpass filter from DC - 280 Hz, DC battery powered by 2 ActiPower PowerPacks (Brain Products GmbH, Germany). Data was recorded continuously with 1 kHz sampling frequency using BrainVision Recorder software (Brain Products GmbH, version 1.21.0303) on an independent recording PC running Microsoft Windows 7. Triggers from the experimenter PC to the recording system were sent via the parallel port.

#### Eye tracking data acquisition

Eye position was recorded using a desktop mount EyeLink 1000 Plus eye tracking system (SR Research Ltd.) and EyeLink 1000 Plus Host software running on an independent recording PC. Connection between the experimenter PC and the EyeLink recording system was established via ethernet, and controlled using the PyLink Python module (SR Research Ltd., version 1.11.0.0) Epoched recordings were made for each trial, with a sampling frequency of 500 Hz. The eye tracker was re-calibrated every other block using a horizontal and vertical calibration routine at 5 positions.

In order to enhance data quality the participants were asked in the email invitation to not apply make-up before the study, especially eyeliner and mascara.

### 2.5 Data preprocessing and processing

Data preprocessing, processing, visualization, and analysis was done using Python (version 3.7), Matlab (The Math-Works Inc., version R2018a), and R (Team, 2018, version 3.5.0).

#### EEG data preprocessing

EEG Data were transferred into BIDS format (C. R. Pernet et al., 2019) using the MNE-BIDS Python module (version 0.4 Appelhoff et al., 2019).

EEG data were only minimally preprocessed with the PREP pipeline (Bigdely-Shamlo et al., 2015), implemented in EEGLAB (Delorme & Makeig, 2004), using default parameters. The pipeline is an automated and fully reproducible multistage preprocessing routine consisting of line noise removal, highpass filtering, robust rereferencing to average reference, detection of noisy channels, and subsequent interpolation of these channels. PREP reports can be found in our online repository. Other than mentioned in the preregistration, we applied no rejection of data based on visual inspection to enhance reproducibility of the study.

To further evaluate the proxy metric, we generated an alternative version of the EEG data more thoroughly cleaned with the AUTOMAGIC preprocessing pipeline (Pedroni et al., 2019). This pipeline combines the early stage preprocessing of the PREP pipeline with a fully automatic ICA based artifact removal (MARA; Winkler et al., 2011) to clean the EEG data from common sources of measurement noise, including eye movements.

#### EEG processing and ASSR detection

The analysis of the auditory steady-state response (ASSR) was performed using MNE-Python (Gramfort, 2013, version 0.22.0) and custom code. All analyses were done in sensor space. 8 s epochs were extracted starting from the beginning of each visual stimulus presentation (the last 4 s of the longer movie trials were omitted to yield the same amount of data across all trial types).

The metric we extracted for the present study is the signal-to-noise ratio (SNR) of the ASSR, a local ratio of the power at the stimulation frequency versus the average power in neighboring frequencies (Meigen & Bach, 1999). For each combination of sensor and trial, power spectral density (PSD) was calculated using FFT on the full 8 s of visual stimulation. The PSD spectrum was then transformed into a spectrum of signal-to-noise ratio (SNR) using custom code: the SNR measure we use is defined as the power in a given frequency band relative to the average power in the neighboring frequency bins. Therefore we convolved the PSD arrays along the frequency axis with the following convolution kernel: 6 surrounding bins (3 on each side), skipping the 2 directly neighboring bins. This yields the SNR spectrum for every trial and channel, composed of unitless values. A SNR-value for a given frequency, channel, and trial can take any positive value (SNR > 0), but in the absence of narrowband rhythmic activity in this frequency should be approximately 1. Values much bigger than 1 indicate narrowband rhythmic components in the EEG, as expected for ASSR (but also for power line noise, if not removed). As the dependent measure of interest for this study we extract only SNR at stimulation frequency, i.e. 40 Hz for every trial and EEG channel. The code created for this task was also made publicly available as part of a tutorial in the MNE-Python documentation.

For the subsequent statistical analysis, the resulting SNR data arrays were averaged in two dimensions: over all channels of the montage resulting in one SNR value per trial per participant, and subsequently over subsets of trials, resulting in one SNR value per stimulus condition per participant. Unlike typical studies working with ASSR we did not confine the analysis to an auditory region of interest (ROI) on the scalp where the responses are strongest, but averaged data from all registered channels; while an ROI analysis would result in higher average SNR values, such a design could only reflect distortions in these channels while spatially localized noise outside the ROI would be neglected.

Although not preregistered, we decided to log transform SNR values to account for the variable’s skewed gamma distribution, which we hadn’t considered in advance (see Supp. 5.3). Transformation took place on the trial level, after averaging SNR values over all EEG channels. After transformation, the distribution indeed assumed a more gaussian shape and this step did not change the significance pattern of the main ANOVA model.

#### Eye tracking data preprocessing and processing

Recorded data consisted of the raw binocular gaze path (as x/y coordinates for both eyes), but also contained eye blink, saccade, and fixation annotations detected online by the Eyelink system’s proprietary algorithms. The recording files were parsed into an accessible format using the proprietary software EDF Converter (SR Research, version 4.0) and a modified version of the ParseEyeLinkAsc module for Python (https://github.com/djangraw/ParseEyeLinkAscFiles, code based on version 7/4/19).

Blink annotations were taken from the recorded files (detection based on SR default algorithm) but custom code was used to select only binocular blinks (overlapping blinks in traces of both eyes), and only blinks longer than 30 ms.

Saccades and microsaccades were detected from the raw gaze coordinates using the velocity based detection algorithm proposed by Engbert and Mergenthaler, implemented in the Microsaccade Toolbox for R (Engbert & Mergenthaler, 2006, version 0.9). Parameters were set to binocular saccades, a minimal saccade duration of 6 ms (3 samples), and a relative velocity threshold (VFAC parameter) of 5. This procedure was preregistered as an optional step to increase data quality. Saccades with a peak velocity ≥ 900°/*s* were rejected as artifacts. Saccades with an amplitude ≤ 1°*deg* were categorized as microsaccades.

To align with the EEG data, timing of blinks and saccades was expressed relative to trial onset (as identical triggers were sent from the presentation PC to both the EEG amplifier and eye tracker.) Blinks and saccades outside the trial limits were discarded. For the statistical analysis, counts per trial for blinks, microsaccades and saccades were computed as dependent trial measures (8s beginning with the onset of the visual stimulus).

#### ECG preprocessing and processing

ECG data handling and initial preprocessing of the unsegmented recording was done in MNE Python, processing and detection of R-peaks and heart rate in trial segments was done using HeartPy module for python (van Gent et al., 2019, version 1.2). The data were visually inspected to detect whether polarity of the recording electrodes was correct, if not this was accounted for during preprocessing.

The initial processing pipeline consisted of a bandpass filter between 0.01 - 100 Hz (FIR filter, zero phase), a 50 Hz notch filter to remove line noise (zero phase), cutting epochs of −8 to 8 s around visual stimulus onset. Heartrate (HR) was then computed for every trial within a baseline window (8 s immediately preceding the visual stimulus onset) and a trial window (8 s starting with visual stimulus onset) by the following pipeline: In each segment the polarity of the recorded ECG signal was switched if necessary, a notch filter was applied to remove baseline drift (0.05 Hz notch, zero phase), and the signal was scaled to have a more or less constant amplitude over time (to help the detection algorithm, HeartPy default parameter). The R-peak was detected from the preprocessed ECG, outlier were detected based on the interquartile range method (IQR): The IQR or midspread is defined as the difference between the 75th and 25th percentiles of the data distribution. Values outside the range of mean ± 1 IQR were regarded as outliers and substituted with the median value. The interbeat intervals of accepted R peaks were averaged and converted to heartrate (HR) in beats per minute (bpm).

For each trial HR values for baseline and trial segments were extracted for all participants. The difference between HR of the baseline segment and HR of the stimulus segment was computed for every trial as the measure of interest. Outlier in HR deceleration values were removed by a conservative, visually derived threshold (lower limit: −30 bpm, upper limit: +30 bpm) that rejected data from 13 trials. We note that this analysis of the ECG data was not preregistered in detail, but only generally listed as an exploratory investigation.

### 2.6 Statistical analyses and visualization

#### ANOVA model

Main effects and interactions of the experimental conditions on the dependent measures were tested using a categorical three-way fully crossed ANOVA with 8 groups with 3 factors in a within subject design (repeated measures ANOVA). The three factors were viewing task (fixation/free), stimulus motion (video/static) and content (dance/landscape). We note that in the preregistration, while we described the intended three-way ANOVA we erroneously wrote about a “two-way” ANOVA; we apologize for any confusion.

This repeated measures ANOVA model was applied to the following dependent measures individually: ASSR SNR (log transformed average over all sensors for each trial, see below and Supp. 5), aesthetic ratings, boredom ratings, blink rate, saccade rate, and microsaccade rate (as counts per trial) and heart rate deceleration (in bpm). Responses were therefore aggregated and averaged within participants for each of the 8 possible factor combinations (i.e. averaging over 10 trials within each participant). Unfortunately a bug in the randomized allocation of the stimuli to the viewing task condition led to a slight imbalance in some of the groups. This error occurred for half of the participants: While overall full balance was kept for the main effects (n=1674-1675 trials each in all participants) and for two of the two-way combinations (stimulus motion x content and fixation task x content; n=834-840 trials in each combination), the combination of fixation task x stimulus motion (n=614-615 trials in dynamic x free gaze and static x fixation, but n=1060 trials in dynamic x fixation and static x free gaze) as well as the three-way combinations were unbalanced (n=530 trials or n=304-310 trials). However, each participant was presented with at least 5 trials per combination (instead of 10 as laid out), and as the ANOVA model was applied to averaged values per participant the effect should be manageable.

The models were created using least squared regression implemented in the AnovaRM class from the statsmodels module for Python (Seabold & Perktold, 2010, version 0.12.1). An effect size (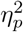: partial eta squared) for all main effects and interactions was calculated from the ANOVA table using the formula for fixed effect designs

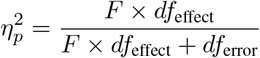

implemented in custom code (after Lakens, 2013). Full ANOVA tables can be found in the supplemental material 5.3).

#### Non-parametric cluster-level paired t-test

To statistically compare the topography of the participants’ ASSR we performed non-parametric cluster-level paired t-tests (Maris & Oostenveld, 2007) for the three main effects (fixation task vs. free-viewing condition, video stimuli vs. still images, dance vs. landscape scenes) using MNE-Python’s permutation_cluster_1samp_test function. MNE-Python’s implementation of this test internally applies a 1-sample t-test on the difference in observations (mathematically equivalent to a paired t-test) and uses sign flipping for generating random permutations. For each participant, log-transformed SNR for each participant and channel was averaged across all trials of the respective main effect categories (fixation task, free-viewing, video stimuli, still images, dance scenes, landscape scenes) and contrasts for all participants were formed by subtraction. Arrays with channel-wise contrasts for all participants were then subjected to the cluster based permutation with 1024 permutations and a t-threshold automatically chosen to correspond to a p-value of 0.05 for the given number of observations. A spatial adjacency matrix for the channel layout was provided based on the easycap-M1 template.

#### Correlation analysis

We analysed the trial wise correlation between several of the response measures: aesthetic and boredom ratings, SNR and both ratings, SNR with eye blink count, saccade count and microsaccade count, and both ratings with eye blink count, saccade count and microsaccade count.

Therefore we applied repeated measures correlation (Bakdash & Marusich, 2017) to take into account that individual observations were clustered by participant; we used the method implemented in the rm_corr function from the pingouin module for Python (Vallat, 2018, version 0.3.8). To construct the rmcorr model of the blink data we had to remove data from 2 additional participants who exhibited 0 detectable blinks during all of the trials (the model did not converge). All correlation results were corrected for multiple comparisons using Holm’s method (Holm, 1979) implemented in the multipletests function from the statsmodels module for Python (Seabold & Perktold, 2010, version 0.12.1). Correlation pattern of all dependent variables can be found in the supplemental material 5.3).

Local regression in Fig. 2c was generated using locally weighted scatterplot smoothing (LOWESS) as implemented in the seaborn module for Python (Waskom, 2021, version 0.11.1)

**Figure 2:**
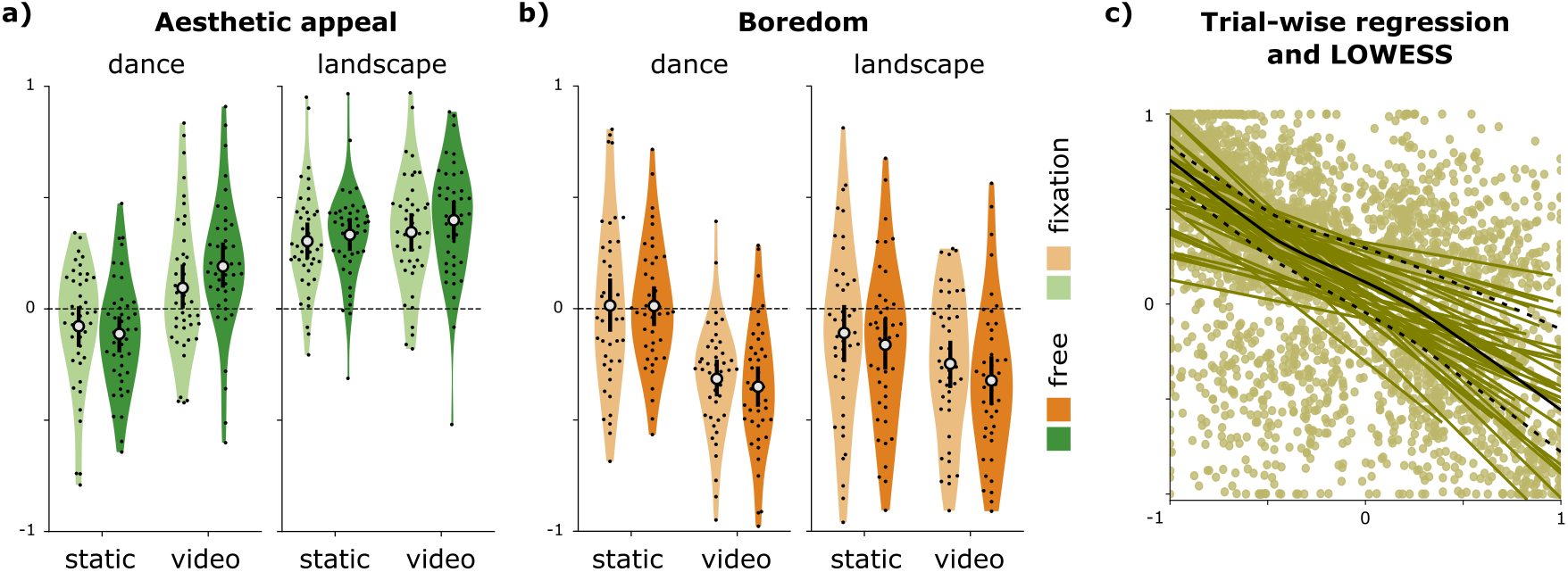
Effect of presentation category on stimulus ratings and correlation of aesthetic appeal and boredom ratings (N=43 participants). a) Average aesthetic rating by category (one dot for each participant): landscapes were rated significantly higher than dance performances (p < .001), dynamic video stimuli were rated significantly higher than static images (p < .001), and stimulus motion (video/static) interacted significantly with content (dance/landscape; p < .001) suggesting that dynamic video stimuli affected the appeal of dance to a greater extent compared to landscapes. b) Average boredom rating by category (one dot for each participant): significantly lower boredom ratings for videos compared to static images (p < .001), and stimulus motion again interacted significantly with stimulus content (p < .001) with the same effect direction. c) Correlation between aesthetic and boredom ratings (one dot for each trial): ratings of aesthetic appeal and boredom were negatively correlated (*r*(3305) = −.58, *p* < .001). This relation was very robust for individual observers (individual lines). A locally weighted fit of all data (LOWESS; solid black line) suggests that the relationship has a degree of nonlinearity and that this nonlinearity is stronger in landscape (upper dotted line) than in dance stimuli (lower dotted line): a larger number of landscape trials were rated as boring yet still moderately aesthetically appealing.

#### Visualization

Data visualizations were generated using the seaborn Python module (Waskom, 2021) with custom post-processing using Inkscape (https://inkscape.org).

## 3 Results

Observers viewed videos and images of natural landscapes and dance performances and were either allowed to freely view the stimuli or asked to fixate a central fixation dot in a fully crossed factorial design (2 stimulus motion x 2 content x 2 tasks) while EEG, eye tracking and ECG data were recorded. Following each trial, observers rated the stimulus for aesthetic appeal and for their state of boredom. Main effects and interactions of the experimental conditions on the various dependent measures were tested using a categorical three-way repeated measures ANOVA with 8 groups and 3 factors: viewing task (fixation/free), stimulus motion (video/static) and content (dance/landscape). Average ratings of the trials within each of the 8 groups per participant were used. Full ANOVA tables can be found in the supplemental material 5.3.

### 3.1 Behavioral measures of aesthetic appeal and boredom

Average ratings of both aesthetic appeal and of boredom were strongly affected by experimental condition (Fig. 2a and b). The overall 3-way ANOVA for aesthetic ratings was highly significant (F(301,42) = 5.87, *p* < .001), accounting for approximately 81% of the variance in participant’s ratings (*adj.R*^2^ = 0.81). The 3-way ANOVA for boredom ratings was also highly significant (*F*(301,42) = 3.84, *p* < .001), and accounted for approximately 71% of variance in aggregate participant ratings (*adj.R*^2^ = 0.71. Both models show several significant main effects and interactions.

As hypothesized, aesthetic ratings of natural landscape stimuli were significantly higher compared to the dance performances (Fig. 2a; *F*(1,42) = 50.21, *p* < .001, 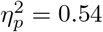). This replicates findings from previous research that nature scenes tend to be rated higher than various other visual stimulus categories (Isik & Vessel, 2019; Vessel et al., 2018).

In addition, dynamic video stimuli were rated as significantly more appealing than static images (*F*(1,42) = 29.72, *p* < .001, 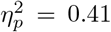) and stimulus motion (video/static) interacted significantly with content (dance/landscape; *F*(1,42) = 18.50, *p* < .001, 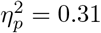). In particular, moving from video to static images affected the appeal of dance to a greater extent than for landscapes. No other main effect or interaction reached significance. However, the effect of fixation task on ratings of aesthetic appeal just slightly exceeded the significance threshold (aesthetic appeal slightly lower under attempted fixation; *F*(1,42) = 4.02, *p* = .051, 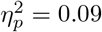).

Turning to the boredom ratings, we observed significantly lower boredom ratings for videos compared to static images (Fig. 2b, *F*(1,42) = 41.18, *p* < .001, 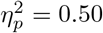), confirming that restricting stimuli to static images has a negative effect on engagement. As for aesthetic ratings, stimulus motion again interacted significantly with stimulus content (dance/landscape; *F*(1,42) = 18.92 *p* < .001, 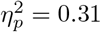), with the shift from video to static images having a greater effect with dance stimuli than with landscape stimuli. Although we had hypothesized that the free-viewing task would result in lower boredom ratings compared to the fixation task, this effect, though in the predicted direction, was not significant (*F*(1,42) = 2.16, *p* = .15, 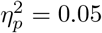). No other main effect or interaction reached significance.

As expected, aesthetic ratings and boredom ratings were negatively correlated (*r*(3305) = −.58, 95*CI* = [−.60, −.56], *p* < .001; Fig. 2c and Supp. Tab. 5). The relationship was extremely robust for individual observers, whom all showed a negative relationship (Fig. 2c, individual lines). A locally weighted fit of all data (LOWESS; solid black line in Fig. 2c) suggests that the relationship has a degree of nonlinearity, being more steep at the extremes but shallower for middle ratings. Separate LOWESS fits for landscape (upper dotted line in Fig. 2c) and dance stimuli (lower dotted line in Fig. 2c) reveal that this might be attributed to a less linear relationship for landscape stimuli compared to dance: a larger number of landscape trials were rated as boring yet still moderately aesthetically appealing.

### 3.2 Physiological measures

#### Eye tracking

In order to better understand how relaxing experimental constraints might affect both behavior and EEG signal, average blink rate, saccade rate, and microsaccade rate were extracted from eye tracking data (aggregated to counts per trial; see Fig. 3).

**Figure 3:**
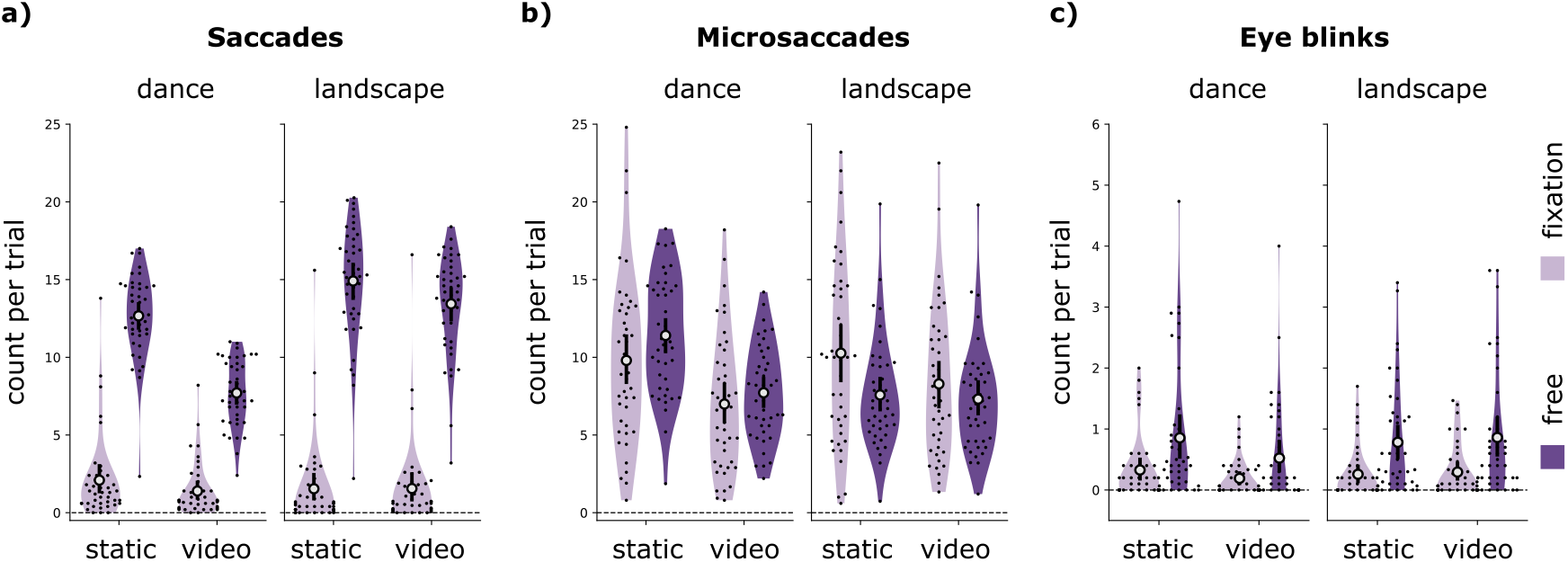
Effect of presentation category on gaze (N=43 participants). a) Larger saccades (> 1°deg) were significantly decreased by fixation task (*p* < .001), but there were also significantly fewer saccades for video stimuli compared to static pictures (*p* < .001), and for dance compared to landscape (*p* < .001). All interaction effects reached significance. b) There were significantly fewer microsaccades (< 1°*deg*) when viewing dynamic videos than when viewing static scenes (*p* < .001), and when viewing landscapes compared to dance (*p* = .025). Further there was a significant interaction between stimulus content and stimulus dynamics, with a more pronounced effect of video in the dance stimuli compared to landscapes (*p* < .001), a significant interaction between stimulus content and viewing task (*p* < .001), and a significant three-way interaction (*p* = .008). c) Eye blink rate was significantly decreased by fixation task (*p* < .001) and video stimuli (*p* = .031), with a significant interaction effect between stimulus content and dynamics (*p* < .001).

Not surprisingly, eye movements were significantly affected by viewing task, but also by stimulus dynamics and content. The ANOVA models for all three measures were significant, though the amount of variance accounted for varied across the three measures. For the number of larger saccades, the model captured 96% of the variance (Fig. 3a, *F*(301,42) = 30.79, *p* < .001, *adj.R*^2^ = 0.96). For microsaccade counts, the model accounted for only 78% of variance (Fig. 3b, *F*(301,42) = 4.93, *p* < .001, *adj.R*^2^ = 0.78), and for eye blink counts, the model accounted for approximately 86% of variance (Fig. 3c, *F*(301,42) = 7.75, *p* < .001, *adj.R*^2^ = 0.86).

Viewing task modulated larger saccades and blink count (both of them were more frequent in the free viewing condition), with a particularly strong effect on larger saccades (*F*(1,42) = 420.17 *p* < .001), 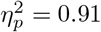; eye blinks: *F*(1,42) = 27.46 *p* < .001, 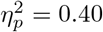). Thus enforcing fixation did indeed significantly reduce eye movements, as intended. Interestingly, however, we observed no significant effect of the fixation task on microsaccade count.

Saccade count was also strongly affected by both stimulus motion and stimulus category, and all interaction effects reached significance. Perhaps counterintuitively, there were more saccades when viewing static images than videos (*F*(1,42) = 152.88 *p* < .001, 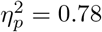), and more saccades for landscape stimuli than for dance stimuli (*F*(1,42) = 135.02, *p* < .001, 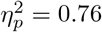). Stimulus motion interacted with stimulus content such that there was a stronger effect of motion for the dance stimuli (*F*(1,42) = 58.36, *p* < .001, 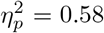), and stimulus motion also interacted with task, with a stronger effect of task for static images (*F*(1,42) = 125.92, *p* < .001, 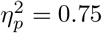). Stimulus content and task also interacted significantly, with a stronger effect of task for landscape stimuli (*F*(1,42) = 113.23 *p* < .001, 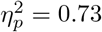). Finally, the three-way interaction was also significant (content x motion x task), with the task by motion interaction being larger for dance stimuli, resulting in the comparatively smallest effect of task for dance videos (*F*(1,42) = 28.85 *p* < .001, 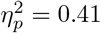).

Interestingly, microsaccade counts were significantly affected by stimulus motion, with fewer microsaccades when viewing dynamic videos than when viewing static scenes (*F*(1,42) = 79.49 *p* < .001, 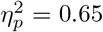), and by stimulus content, with more microsaccades in dance than in landscape (*F*(1,42) = 5.42 *p* = .025, 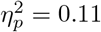). There was also a significant interaction between stimulus content and viewing task, with a differently oriented effect of fixation task for the two content conditions (fewer microsaccades under fixation in dance, more so in landscapes; *F*(1,42) = 31.25 *p* < .001, 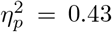). Further there was again a significant interaction between stimulus content and stimulus motion, with a stronger effect of motion on number of microsaccades for the dance stimuli compared to landscapes (*F*(1,42) = 24.13 *p* < .001, 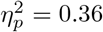). The three-way interaction was also significant (*F*(1,42) = 7.83 p = .008, 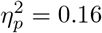).

Besides the effect of viewing task, eye blinks were marginally significantly affected by stimulus motion, with fewer blinks during dynamic video clips (*F*(1,42) = 4.96 p = .031, 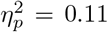), and a significant interaction between stimulus dynamics and stimulus content (*F*(1,42) = 9.89 p = .003, 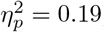).

We observed several significant trialwise correlations between eye movements and the behavioral ratings: both aesthetic ratings and boredom ratings were significantly correlated with eye blink count (aesthetic appeal: *r*(3305) = −.06, 95*CI* = [−.09,−.02], *p* < .001; boredom: *r*(3305) = .08, 95*CI* = [.04, .11], *p* < .001) and microsaccade count (aesthetic appeal: *r*(3305) = −0.09, 95*CI* = [−.12, −.05], *p* < .001; boredom: *r*(3305) = .13, 95*CI* = [.09, .16], *p* < .001), with boredom exhibiting a slightly stronger correlation. Saccade count did not significantly correlate with the ratings.

#### ASSR SNR

One of our primary goals was to assess the effects of task and stimulus motion on EEG signal quality. To do so, a continuous auditory stimulus (pink noise with 40 Hz amplitude modulation; see Methods) was played during each trial, and SNR of the auditory steady-state response (ASSR) was computed as a proxy measure for overall EEG recording quality.

Despite their strong effects on behavioral ratings of aesthetic appeal and boredom, the experimental manipulations had only minimal effects on measured SNR (Fig. 4a). Average SNR values were slightly higher under attempted fixation compared to free viewing. The overall ANOVA model was significant (*F*(301,42) = 8.57, *p* < .001, *adj.R*^2^ = 0.87), with fixation task as the only significant factor (*F*(1,42) = 6.24 p = .017, 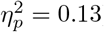).

**Figure 4:**
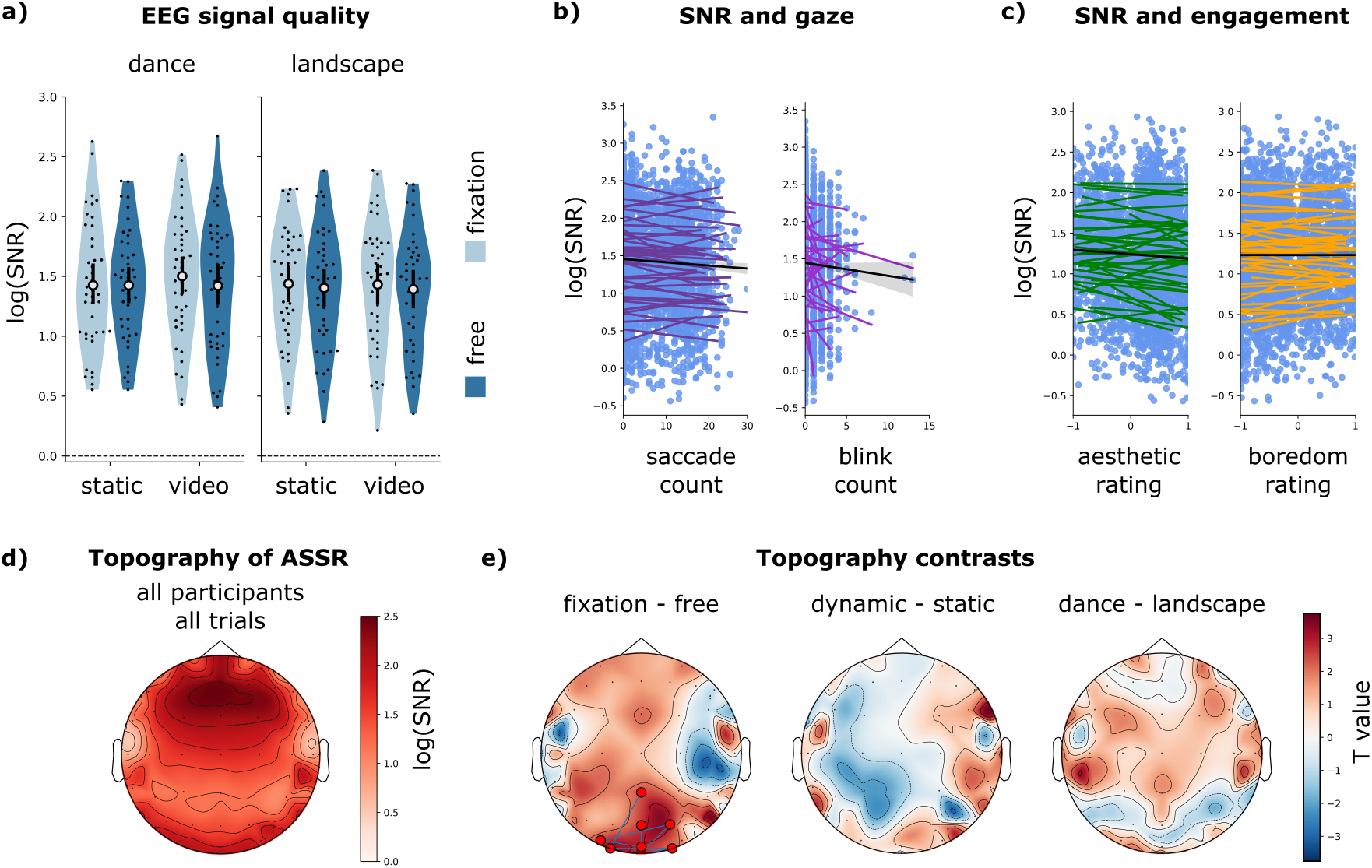
Effect of presentation category on ASSR SNR and correlation of SNR with saccade count, eye blink count, and behavioral ratings (N=43 participants). a) Significantly higher SNR for fixation task compared to free gaze (p = .017). b) Significant trialwise correlation (repeated measures correlation) of log transformed SNR with the participants’ number of eye blinks per trial (*r*(3305) = −.05, *p* = .04) but not with number of saccades per trial (*r*(3305) = −.04, *p* = .18). c) No significant trialwise correlation (repeated measures correlation) between log transformed SNR and aesthetic ratings (*r*(3305) = −.05, *p* = .10), and marginally significant correlation between log transformed SNR and boredom ratings (*r*(3305) = .04, *p* = .26). d) ASSR is detected at all electrode locations, with a maximum over fronto-central vertex channels (common average reference; grand average log-transformed SNR across all participants and trials). e) Overall significant differences between fixation task and free viewing condition - see panel a) - are mainly caused by occipital channels (Nonparametric cluster-level paired t-test; significant channels with *correctedp* < .05 marked in red). No significant differences in any channel or cluster detected for video vs. still images, or dance vs. landscape stimuli.

Importantly, while we found only a small effect of the experimental conditions on SNR values, we believe that the measure is sufficiently sensitive to adequately capture bad quality data and noisy trials. The trialwise SNR measure did correlate with the number of blinks per trial (*r*(3305) = −.05, 95*CI* = [−.09, −.02], *p* = .002) and with the number of saccades (*r*(3305) = −.04, 95*CI* = [−.08, −.01], *p* < .018; see Fig. 4b). However, only the correlation with blink count survived the multiple comparisons correction (*p.corr* = .036). The SNR measure did not correlate with the number of microsaccades per trial (repeated-measures correlation *r*(3305) = .00, 95*CI* = [−.03, .03], *p* = .98).

When directly compared with the behavioral ratings (Fig. 4c), there was a trialwise correlation between log transformed SNR and boredom ratings (*r*(3305) = .04, 95*CI* = [.00, .07], *p* = .029) as well as with aesthetic ratings (*r*(3305) = −.05, 95*CI* = [−.08, −.01], *p* = .008); however these correlations were not strong enough to remain significant after multiple comparisons correction (see Supp. Tab. 5 for full trialwise correlation structure).

Fig. 4d) shows the average topography of the ASSR on the scalp (grand average over all subjects and all trials per electrode): the measured ASSR reached its maximum over fronto-central vertex channels, congruent with previous work using EEG data with common average re-referencing (e.g. Lustenberger et al., 2018). Additionally we see potential dipole patterns over left and right temporal cortex that might reflect a generating source of the ASSR component in auditory brain regions.

We statistically contrasted the topographies of the ASSR for the different investigated presentation conditions to see whether there were any region specific effects not seen in the analysis of SNR values averaged over the full scalp (see Fig. 4e). A nonparametric cluster-level paired t-test shows that the fixation task significantly enhanced ASSR SNR solely in occipital channels. Using video stimuli rather than still images, as well as the content condition (dance vs landscape scenes) did not result in significant differences in any channel or channel cluster, which is congruent with the absence of any overall significant effect of these conditions in the ANOVA models.

To further evaluate the proxy metric, we repeated the entire analysis on EEG data that were cleaned in a fully automated preprocessing pipeline with ICA based artifact removal (AUTOMAGIC Pedroni et al., 2019). After this step there was now no effect of any of the investigated factors on log transformed SNR: the overall ANOVA model was significant (*F*(301,42) = 7.74, *p* < .001, *adj.R*^2^ = 0.86) but none of the main effects nor interactions were significant (effect of fixation vs. free gaze *F*(1,42) = 0.67 *p* = .42), 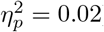). Unlike in the minimally preprocessed data, log transformed SNR in the cleaned data did not significantly correlate with saccade count (*r*(3305) = −.02, 95*CI* = [−.05, .01], *p* = .24), blink count (*r*(3305) = −.02, 95*CI* = [-.06, .01], *p* = .18) nor with microsaccade count (repeated-measures correlation *r*(3305) = .02, 95*CI* = [-.02, .05], *p* = .35).

Importantly, this is not due to the fact that the ASSR component was removed from the EEG in this preprocessing step: the AUTOMAGIC routine applies a pretrained ICA classifier (MARA Winkler et al., 2011) that is not expected to remove auditory brain components. While log transformed SNR values in the ICA cleaned data were, to our surprise, significantly lower than in the minimally preprocessed data (grand average 1.27 vs. 1.43; 2-sided paired *t* = −10.38, *p* < .001), they were still significantly larger than 0 (equivalent to a raw SNR=1; 1 sample *t* = 46.00, *p* < .001).

#### ECG

We performed an exploratory analysis of event-related heart rate (HR) changes to investigate the participants’ autonomous responses to the stimuli. We observed a significant HR deceleration after the onset of the visual stimuli, consistent with previous reports (e.g. for IAPS images Palomba et al., 1997). HR during the 8 s time window preceding onset of the visual stimulus (baseline) was significantly larger than HR in the 8 s window after stimulus onset (2-sided paired *t* = 33.00, p < .001, for averaged HR values). However, the investigated experimental factors did not strongly influence HR deceleration. The overall ANOVA model was significant but accounted for only 33% of the variance (*F*(301,42) = 1.55, *p* = .043, *adj.R*^2^ = 0.33). The only significant main effect was stimulus motion (*F*(1,42) = 11.24 *p* = .002, 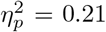), indicating that HR deceleration was slightly stronger for videos than for static pictures.

We further observed no significant correlation of HR deceleration with either aesthetic ratings ( *r*(3292) = .01, 95*CI* = [-.03, .04], *p* = .76) or boredom ratings (*r*(3292) = -.01, 95*CI* = [-.04, .02], *p* = .57). Yet, interestingly HR deceleration significantly correlated with blink rate (*r*(3292) = 0.05, 95*CI* = [0.01,0.08], *p* = 0.005), saccade rate (*r*(3292) = .06, 95*CI* = [0.02,0.09], *p* < 0.001), and SNR (*r*(3292) = -.04, 95*CI* = [−0.07, −0.00], *p* = 0.039), but not with microsaccade count (*r*(3292) = −.00, 95*CI* = [-0.03,0.03], p = 0.98). Only the correlation with saccade count was strong enough to remain significant after multiple comparisons correction (*p.corr* = .012; see Supp. Tab. 5 for full trial wise correlation structure).

## 4 Discussion

In the present study we assessed trade-offs between ecological validity and EEG signal quality as participants viewed complex, aesthetically engaging stimuli. We found that the use of video stimuli does not necessarily result in lower EEG quality but can in fact significantly reduce eye movements under free viewing conditions, especially if human agents are depicted (see Table 1 for a comprehensive overview of the main effects). The use of video stimuli further resulted in significantly higher aesthetic ratings and lower perceived boredom, indicating higher engagement with the task and stimuli. This constitutes another benefit of using video material, given that a lack of engagement might in turn decrease data quality or even cause untimely termination of data collection. The fixation task was confirmed to significantly reduce eye movements and to have an overall positive effect on EEG quality. The effect on SNR was mainly driven by differences in occipital channels. Small fixational eye movements, on the other hand, were not inhibited by attempted fixation. Though we observed a trend for lower aesthetic ratings under attempted fixation, the fixation task did not significantly influence the investigated behavioral ratings. This is encouraging given the predominance of this experimental constraint in studies using EEG and MEG. Finally, the stimulus content domain, primarily included as a control condition for the behavioral responses, had a remarkably far reaching influence. Beyond its already known effect on aesthetic ratings it also significantly affected saccade count and microsaccade count, and showed consistent interaction effects with the stimulus dynamics in several of the behavioral measures. By presenting these data and findings we hope to help inform future trade-offs during the design and piloting phase of neuroimaging experiments and to encourage researchers to reconsider the default application of canonical constraints.

**Table 1:**
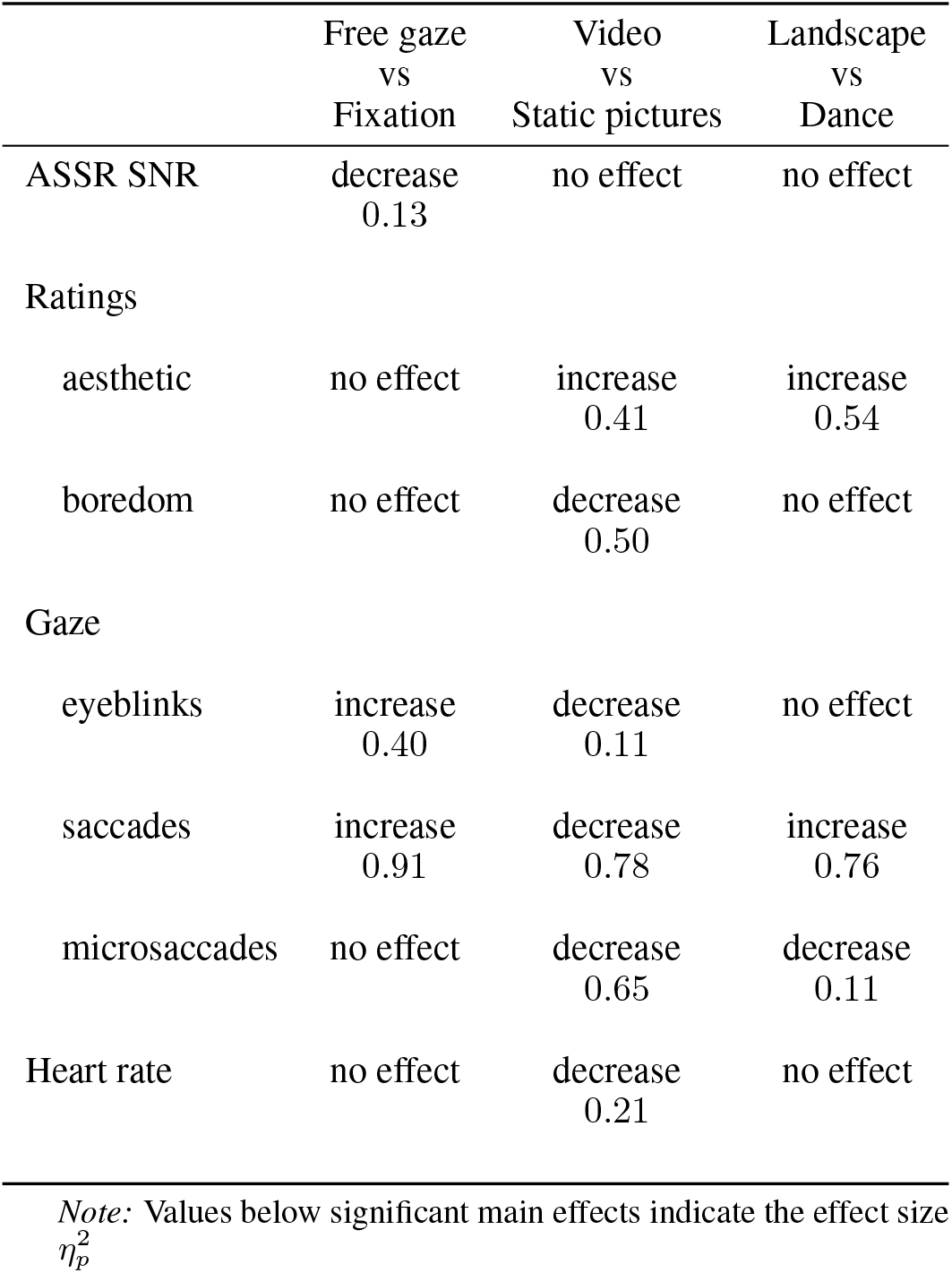
Main effects of experimental conditions on dependent variables

|  | Free gaze<br>vs<br>Fixation | Video<br>vs<br>Static pictures | Landscape<br>vs<br>Dance |
| --- | --- | --- | --- |
| ASSR SNR | decrease<br>0.13 | no effect | no effect |
| Ratings |  |  |  |
| aesthetic | no effect | increase<br>0.41 | increase<br>0.54 |
| boredom | no effect | decrease<br>0.50 | no effect |
| Gaze |  |  |  |
| eyeblinks | increase<br>0.40 | decrease<br>0.11 | no effect |
| saccades | increase<br>0.91 | decrease<br>0.78 | increase<br>0.76 |
| microsaccades | no effect | decrease<br>0.65 | decrease<br>0.11 |
| Heart rate | no effect | decrease<br>0.21 | no effect |

*Note:* Values below significant main effects indicate the effect size $\eta_p^2$

### 4.1 Using ASSR SNR as a proxy measure for EEG signal quality

We had hypothesized that without extensive preprocessing of the EEG data SNR of the auditory steady-state response would be affected by the viewing task (i.e. lower SNR in free-viewing condition than in the fixation task). Indeed, we observed that the fixation task enhanced grand average SNR across all conditions. The same effect was present in eye blinks and saccades (significantly fewer of these with fixation task) and SNR was significantly correlated with the number of blinks on a trial-by-trial basis. In order to evaluate the reliability of the proposed proxy measure and to look for significant effects of the presentation conditions we did not optimize the preprocessing pipeline for highest possible SNR response. We applied only minimal frequency domain filtering, and most importantly we did not exclude or reconstruct “noisy” data segments (except for rejection and interpolation of outlier channels): for the primary analysis we applied no visual inspection of trials, nor any ICA or EOG based cleaning. Furthermore we averaged over the full scalp montage rather than using an ROI based analysis (i.e. no spatial cleaning that would differentially remove or reduce noise components with certain sources or orientations).

Any of these cleaning approaches would likely result in higher SNR measures and a better detection of ASSR. For example, as the topography of the ASSR (Fig. 4d) shows, confining analysis to a fronto-central ROI would have resulted in higher average SNR values. However, analyzing only data from a fronto-central ROI we would likely have missed the effect of the fixation task, as it was mainly driven by SNR differences in occipital channels (see Fig. 4e). Keep in mind that in our framework the ASSR generated by the brain is considered a fixed factor, while we are interested in detecting *changes in ASSR SNR* measured at the scalp that are caused by different types of noise. Thorough cleaning of the data would thus be counterproductive, as it removes the negative effects of noise that we are interested in quantifying. This was confirmed by our secondary analysis of a version of the EEG data cleaned via an ICA-based artifact rejection pipeline. In these data, the significant effect of the fixation task vanished, as did the correlation of the SNR measure with saccade count and blink count.

This leads us to the conclusion that the SNR measure used in this study indeed captured eye movement related signal distortions that would typically lead to a rejection of trials or data segments in common EEG cleaning routines (Luck, 2014). As Fig. 4d shows, ASSR signal was measured on all scalp electrodes and hence noise distortions on any channel could have lowered the grand average SNR used as a dependant variable in this study. However, the effect size associated with the fixation task was small and there were large differences across participants in their average SNR, indicating that that the SNR values may be meaningfully compared only within one subject, but not across participants. Hence, even though ASSR SNR shows some sensitivity and can capture noise in the EEG, its promise as a universal quantitative metric for EEG quality is questionable. We will discuss this point in more detail below.

### 4.2 Using video stimuli improves engagement, reduces eye movements, and only minimally affects EEG SNR

Importantly, we found evidence for our hypothesis that the experimental restrictions compared in this study not only affected EEG signal quality, as intended, but also several other dependent measures. We observed higher aesthetic and lower boredom ratings for videos than for static stimuli. While the two measures might focus on slightly different psychological evaluation processes, both likely reflect an engagement component and hence this finding suggests that video stimuli were experienced as more engaging than the static images generated from the exact same visual content.

While high engagement with stimulus material is certainly important for fields that directly study it, such as empirical aesthetics, we want to emphasize that other fields might also want to design their experiments to be as engaging as possible. Soft factors such as a participant’s commitment, their motivational state, or the engagement with the task can have an influence on data quality, in the worst case leading to early termination of recording sessions. Another possible mechanism by which EEG quality might be affected by a participant’s state of engagement is physical restlessness, which can lead to more frequent body- (muscle artifacts) and eye movements (Ramos Gameiro et al., 2017). The same holds true for fatigue and drowsiness, which impair a participant’s ability to concentrate. In the EEG, however, such effects are to date very difficult to quantify: a growing body of applied research tries to detect states of drowsiness based on the human EEG (Stancin et al., 2021), and the extreme case of sleep proper (marked by sleep spindles and other well characterized sleep stages) can be reliably detected by field experts (e.g. Lacourse et al., 2020). However, this is not a part of typical preprocessing routines in cognitive neuroscience. If anything, bad quality data due to a supposed lack of engagement with the task can be identified based on behavioral performance (e.g. response accuracies, false response rates, or response times). However, in order to do so researchers need prior knowledge to set meaningful performance thresholds, which risks exclusion of participants who are engaged but simply bad at the specific task (thereby potentially introducing bias to their population and data by rejecting lower tails of a distribution). Importantly, cleaning data based on participants’ responses is less straightforward for exploratory studies and tasks that do not have “correct” answers, such as ratings of preference.

In light of the present findings, we want to point out that EEG signal quality (and especially eye movement related artefacts) should not be a major concern when deciding whether to use videos or static images. On the contrary, using video stimuli might even come with further practical advantages besides higher engagement: with video material we observed significantly fewer blinks, microsaccades and saccades, and our proxy measure for EEG quality was not affected by stimulus dynamics. Interestingly, the boredom ratings were significantly correlated with the participants’ eyeblink rate and especially their microsaccade rate, indicating that boredom can increase these potentially problematic eye movements, even despite a fixation task.

Hence, dynamic stimulus material might in fact be a promising tool to naturally reduce eye movements and blinks without having to apply a fixation task with all its potentially negative side effects (see above). Our data suggests that this effect might be especially strong when humans are depicted. These positive effects of video might be related to a phenomenon called center bias (a tendency to shift and keep the gaze focused at the center of a visual stimulus) that appears to be stronger in videos than in static stimuli (for a review see T. J. Smith, 2013). Dynamically changing visual stimuli can also exert some level of exogenous control on observers’ gaze, thereby creating temporal attentional synchrony across viewers (Goldstein et al., 2007; T. Smith & Henderson, 2010; T. J. Smith & Mital, 2013). This latter point further suggests that video stimuli might be used to align both the sensory input stream and endogenous eye-related noise (Nikolaev et al., 2016) across participants, potentially with a similar efficiency as a fixation task. Indeed, hyperscanning studies frequently use video stimuli to investigate similarity of brain responses across observers (Hasson & Frith, 2016; Poulsen et al., 2017).

In current practice, though, most researchers would refrain from using video stimuli, not only in electrophysiological work but in cognitive science in general. There are several reasons for this level of caution, mainly the reductionist view that in many cases stimulus dynamics add unnecessary degrees of freedom that can be taken out of the equation by using static pictures. However, while initial studies suggest that some findings from the huge body of EEG work with static stimuli might translate to dynamic video stimuli (e.g. the N170 ERP component was observed using video material, Johnston et al., 2015) a direct transfer of findings between stimulus modalities needs to be established for each component. As previous work from the eye movement literature and the present work suggest, there seem to be significant differences between static and dynamic visual stimuli both in observers’ visual exploration and gaze patterns (leading to different low-level input streams to the visual cortex) as well as in their higher level cognitive processing (manifesting in differences in preference and boredom ratings).

Researchers from fields that rely on unconstrained vision have developed tools and experimental paradigms that allow for analysis of electrophysiological data recorded under such uncommon conditions (e.g. Ayrolles et al., 2021; Ehinger & Dimigen, 2019; Lu & Ku, 2020). Indeed, we would suggest that a growing literature of neuroimaging studies using dynamically changing visual stimuli, regardless of the specific topic at hand, would be beneficial to the broader field. To date, such work is underrepresented in the neuroscientific literature and could eventually bridge the gap to the body of behavioral work concerned with unconstrained vision.

### 4.3 Fixation task improves EEG SNR, and does not significantly impact ratings, but does introduce cognitive load

The effect of viewing task on SNR and eye movements was expected, as it is the very reason to apply this restriction in the first place. Interestingly, the effect on ASSR SNR seems to be mainly driven by differences in occipital channels (see Fig. 4e, left). This is surprising, as these channels are far away from the expected sources of both the ASSR component (temporal regions over auditory cortex, as indicated by the prominent dipole in Fig. 4 d) and previous work (e.g. Picton, 2011)), as well as of the different noise components caused by eye movements (most prominent on frontal channels; Ball et al., 2009; Plöchl et al., 2012). This localization also does not fit the main projection of the ASSR component onto fronto-central vertex channels.

We confirmed that there were significantly fewer blinks and saccades in trials with a fixation task compared to free gaze trials. Notably, the number of blinks was overall very low, with less than one blink per trial averaged over all participants (0.3 per 8s trial for fixation, 0.8 per 8s trial for free gaze). This is well below spontaneous blink rates reported in the respective literature, even for the free gaze condition (e.g. Jongkees & Colzato, 2016, 7.2 - 18.9 blinks per minute (equivalent to 0.96 - 2.52 per 8 s) in the reviewed cognitive research studies, and 6 - 34.4 blinks per minute (equivalent to 0.8 - 4.59 per 8 s) over a broad variety of tasks and situations in healthy populations). As is to be expected, the number of larger voluntary saccades was heavily reduced as well, from a bit less than one per second in the free viewing to less than one saccade per trial in the fixation task (1.6 per 8s trial for fixation, 12.2 per 8s trial for free gaze). This replicates findings from the eye-tracking literature (e.g. Otero-Millan et al., 2008). Again, the numbers observed in this study were very low compared to spontaneous saccade rates. One reason might be that the stimuli were presented in a relatively small format (< 20° *deg* of visual angle) and could be almost fully captured by near peripheral vision (i.e. without saccadic scanning of the stimulus). We want to emphasize that, as seen in our data and reported before (see e.g. Otero-Millan et al., 2008; Thielen et al., 2019), a fixation task is not suitable to suppress microsaccades. In the present study the number of microsaccades was even slightly higher under attempted fixation (8.8 per 8s trial for fixation, 8.5 per 8s trial for free gaze), though the effect was not significant. The overall microsaccade frequency observed in this study lies at the lower end of an expected range of 1-2 Hz reported in the literature (Engbert & Mergenthaler, 2006). Microsaccades can induce transient low gamma band activity in the EEG that can be misinterpreted as brain activity (Yuval-Greenberg et al., 2008), and hence constitute a source of endogenous noise that can not be counteracted with a fixation task. Unfortunately, cleaning the signal using ICA does not reliably remove the effects of microsaccades from the EEG (Craddock et al., 2016; Dimigen, 2020). If this frequency range of the EEG signal matters, one would hence need to coregister microsaccades to fully account for their effects.

Even more important than the absolute reduction of eye movements might be that the fixation task also reduced the variance of eye movements across conditions (striking differences in saccade rate across stimulus conditions under free viewing, see Fig. 3). Systematic differences in eye movements across conditions can constitute confounds and might lead to false or misattributed findings in downstream analyses (Quax et al., 2019; Thielen et al., 2019).

While our proxy metric showed an overall significant effect of eye movements on EEG quality, it might be that this effect is frequency dependent. To test for this, we performed an additional analysis correlating the oscillatory dynamics of the EEG with eye movements in different frequency bands (see Supp. 5.3). The results show that in lower frequencies (delta and theta) the two signals correlate very strongly (delta: *r*(*horizontal*) = .68, *r*(*vertical*) = .61; theta: *r*(*horizontal*) = .51, *r*(*vertical*) = .47), which probably reflects the large signal offsets induced by rotation of the eyes’ dipoles (offset increases linearly with the size of the eye movement; Plöchl et al., 2012). This finding is of particular relevance for research interested in oscillatory dynamics in these low frequencies: for such studies, removing the fixation task and refraining from rejecting trials with eye movements might not be an option, as it bears the risk of misinterpreting artifacts induced by eye movements as neuronal effects. Small correlations in the alpha (*r*(*horizontal*) = .10, *r*(*vertical*) = .10) and beta band (*r*(*horizontal*) = .03, *r*(*vertical*) = .03) might be less of a concern for future research. There might be some amount of eye induced signal reflected in this frequency range, but it seems unlikely that this might flaw an entire study, especially if approaches like trial averaging or signal cleaning are used. High frequency EEG artifacts in the gamma band, such as those induced by saccadic spike potentials or microsaccades (see Plöchl et al., 2012; Yuval-Greenberg et al., 2008), were not reflected in correlation between the two signals (low gamma: *r*(*horizontal*) = .01, *r*(*vertical*) = .01; high gamma: *r*(*horizontal*) = .01, *r*(*vertical*) = .01), despite the fact that there must have been eye movements during each single trial, regardless of the condition (see Sec. 3 eye tracking). The GFP correlation method seems to be insensitive to these high frequency eye movement artifacts.

Apart from eye movements (and SNR) the fixation task did not significantly affect any of the other dependent measures. On first sight this is a very encouraging finding, given how ubiquitously the fixation task is applied in EEG and MEG studies regardless of the investigated questions. On the downside, the predominance of the fixation task in EEG and MEG research makes it difficult to compare findings with the large body of eye movement literature investigating unconstrained gaze patterns (for a review see Eckstein et al., 2017) or spontaneous blink rates (e.g. Jongkees & Colzato, 2016; Stern et al., 1994), as well as to research done using fMRI or other modalities. More importantly, initial research suggests that there might be significant differences in frequently investigated neuronal processing components under free viewing compared to attempted fixation (s.a. in the N170 Auerbach-Asch et al., 2020).

Last but not least, attempting to maintain fixation over a long period adds atypical cognitive load for a study participant and can lead to fatigue. This is problematic for some high-level cognitive tasks, such as research on aesthetic processing (Brielmann & Pelli, 2017). In fact, aesthetic ratings in our dataset were slightly higher in the free gaze condition than under attempted fixation and this effect just slightly missed the arbitrary p-value threshold of significance (*p* = .0513). Stimuli in the present study were small and could be almost entirely captured using the central field of view. It is possible that this made fixation easier or less intrusive for the participants: there were anecdotal reports in the debriefing that it was especially hard, or even frustrating, to fixate if there were salient features one would like to explore right outside the focus of visual attention. Small stimuli, by design, lower the likelihood that such events will happen, and hence, it is possible that we would have found a larger effect if the stimuli were larger. Additional research is needed to clarify whether the trend we observed may thus actually hint at an existing effect on behavior. It is also possible that the dependent measures we collected (boredom and aesthetic appeal) simply did not capture the psychological state negatively affected by a fixation task: perhaps an additional direct question targeting fatigue, stress, or annoyance would have shown effects. Either way, psychological factors like the ones discussed above can decrease a participant’s concentration on or engagement with the primary task, thereby potentially introducing a new source of noise to the various dependent measures within an experiment. These sources might prove more difficult to track and remove than the eye-related artifacts that were to be avoided in the first place. Future research is needed to further characterize the potential effects of eye movement constraints in the cognitive sciences.

### 4.4 Stimulus content also matters, and dance stimuli are differentially affected by video condition

As expected, landscape stimuli were on average rated more aesthetically pleasing than dance scenes, consistent with previous findings (Vessel et al., 2018; Vessel & Rubin, 2010). Interestingly, this main effect was not significant in the boredom ratings, while overall these two types of ratings showed a fairly high negative correlation.

Looking at the LOWESS curves in figure 2c might help to understand the relation between the two measures. We see a larger fraction of landscape trials with high rated boredom that were nevertheless rated highly aesthetically appealing. This suggests that the aesthetic evaluation of landscapes is guided by a partly different set of features than the evaluation of dance scenes: comparably boring landscapes with less salient features (e.g. meadows compared to a rocky mountain vista) might nevertheless be appealing for their expected affordances (e.g. security, lush vegetation), personal relevance (e.g. because they resemble the region an observer grew up in) or other factors. In dance the correlation of aesthetics and boredom was tighter, potentially indicating that the engagement component is more relevant in this domain.

While we cannot offer a conclusive explanation for this phenomenon, the divergent findings are also promising: they posit a proof of concept that the two behavioral rating scales – for boredom and aesthetic appeal – were not used identically by the participants. Further work will be necessary to investigate the relationship between these measures.

One salient finding in our data is that we consistently observed interaction effects between stimulus content (dance/landscape) and dynamics (video/static). We observed this interaction in behavioral ratings, blink count, saccade counts, and microsaccade counts. The interaction effect always went in the same direction, indicating that the effect of stimulus dynamics was stronger in dance scenes and weaker in landscapes (or that the effect of content was stronger with video and weaker with stills, respectively). As mentioned above, the correlation between aesthetic and boredom ratings might also be affected by the content conditions, with a weaker correlation and stronger nonlinearity in landscape scenes.

One potential explanation for this differential effect of content could be that motion matters more when human agents are depicted (at least compared to landscapes). Again, affordance theory (Greeno, 1994) might offer an interpretation: a landscape can offer a whole set of affordances like landscape features, relevant objects, or humans, plants, and animals within it that one can interact with. These can be hidden or dispersed within the landscape and motion would be only one cue among others to identify these (e.g. if movement is happening in a small part of the scene caused by a small animal that one was not aware of). More likely though, affordances will be identified during active visual exploration. This would fit with our observation that there are significantly more saccades for landscapes than for dance stimuli.

Direct interactions with another human, on the other hand, are arguably largely concerned with communication and understanding of the other’s intentions (e.g. non-verbal communication cues can convey intentions like sympathy or threat). This might also be implicitly happening while observing dance scenes. In our study’s setup, vision is the only sensory modality available to the participant and hence the dancer’s motion might constitute the most important stream of information besides overall outer appearance.

### 4.5 Potentially problematic effects of other experimental conditions in the present study

Here we present a study aiming to assess potential problematic effects of inherent secondary manipulations on primary measures of interest; hence we should also consider whether any secondary experimental manipulations we ourselves introduced might have such influence.

One harsh restriction compared to true natural conditions was the head rest. Without this restriction we would expect increased EEG noise due to muscle movements and physical motion of the cables and sensors, but importantly also brain activity related to the motion and repositioning of the head (Gramann et al., 2021). On the other hand, being asked to hold the head still over a long period might result in fatigue or stiffening neck muscles, thereby introducing noise in the EEG and potentially also lowering engagement or aesthetic appeal. Future research could let participants hold their head freely and record the head position and movements via accelerometer or neck EMG to quantify these factors.

The second and arguably most critical manipulation introduced in this study was the background ASSR stimulus. Unlike the fixation task and stimulus dynamics, the auditory background task was not balanced by any number of trials without auditory stimulation. We therefore cannot directly test for its effect on any of our dependent variables. However, measures were taken to make the stimulus as unobtrusive as possible: the sound stimulus was selected from a set of different candidates in a pilot study (data not shown), and loudness was adapted to the individual sensory threshold of each participant. However, there were individual differences in how the participants perceived the stimulus qualitatively. During debriefing several participants mentioned that they had completely stopped noticing the sound after a while. In contrast, a few other participants mentioned that they had sometimes struggled with a decision whether to incorporate the sound into the summary rating of aesthetic appeal; even though they were aware that the sound was a background manipulation, it was nevertheless experienced as part of an audiovisual aesthetic stimulus. Two comments even pointed at another possible instance of an interaction effect with stimulus content, reporting that they had interpreted the auditory stream as the sound of a helicopter in the landscape videos recorded using drone shots. The auditory stimulus used in this study is pink noise with changing loudness, and individual differences in the qualitative perception of audio noise have been reported (Bergamasco et al., 1976). Concerning the brains’ internal processing of the stimulus, we again emphasize that ASSR is primarily an early auditory response (Picton et al., 2003) and has been shown to not interfere with some types of early visual processing (Keitel et al., 2013). Relatedly, traffic noise and white noise have been shown to reduce sensitivity to other auditory stimuli (in this case verbo-acoustic communication), but do not affect sensitivity in the visual domain (Bergamasco et al., 1976). These findings suggest that the ASSR manipulation might not be critical for studies concerned with visual processes. Studies investigating other auditory processing, however, might be more affected, and bottom-up interactions with higher level cognitive processes can not be excluded at this point. We take it as a positive sign, though, that several people did entirely stop noticing the sound.

### 4.6 Future development of ASSR as a general time-varying SNR measure

With the present work we utilized ASSR as an online marker for EEG recording quality on a trialwise and potentially continuous basis (e.g. combined with a sliding window function). To our knowledge, no universally accepted marker exists and this method could be promising for applied research (e.g. BCI context) or research using mobile EEG when the external noise level cannot be sufficiently controlled. In light of the present results, however, we would urge caution.

While the measure was sensitive to common sources of endogenous measurement noise, namely eye movements, we want to raise some concerns: first, the sensitivity of ASSR for these events was low. Effect sizes and correlation coefficients with the metric were very small (especially in comparison to individual differences across subjects). Related to this latter point, the proxy appeared to be only meaningful within, but not across observers. And lastly, the temporal sensitivity of the ASSR metric is not well characterized and probably limited. Here we used data from 8 s trials; while this might be enough for an online measure in many applied contexts (e.g. in a moving window approach for brain-computer interfaces), it is likely too long for many trial based research applications. Other work investigating ASSR typically analyzes sweeps of 30s and above. This makes sense, because the longer a segment of analyzed data the more cycles of ASSR are represented and the SNR (i.e. the detectability of an oscillatory response in inherently noisy data) should increase linearly. However, this also means that the more data is analyzed, the smaller the relative contribution of a time limited noise event, which might not anymore lead to a significant drop of SNR. In the other direction, it is unclear how short analyzed data segments can get before an ASSR is not detectable anymore even in the absence of prominent measurement noise. Extensive further research would be necessary to validate the applicability and sensitivity of ASSR as an online or trialwise marker for EEG data quality.

Even though we systematically lifted some experimental constraints, the study was nevertheless conducted in a very controlled lab environment. Hence we cannot say much about how the metric would react to typical sources of exogenous measurement noise such as electronic equipment or concussion of the electrodes. Likewise we observed overall very low rates of endogenous noise sources such as eye and body movements (the head was mounted on a chin rest), or concurrent sensory input (the experiment took place in an acoustically and visually shielded cabin). These conditions fundamentally differ from the largely uncontrollable recording environments in applied and mobile EEG studies that would most profit from a validated online measure of signal quality.

Artifactual oscillatory activity in the low gamma-band, caused by microsaccades (Yuval-Greenberg et al., 2008), might be problematic for measuring SNR of the 40 Hz ASSR in our study since microsaccade rate was significantly linked to several of the investigated stimulus categories. Also saccadic spike potentials can generate broadband high frequent distortions (Plöchl et al., 2012). It seems possible that ASSR SNR mainly reacts to these high frequent noise components, as they can be in the same frequency range as the signal (40 Hz), even though less confined. If this were true, it would question the usability of the proxy metric. However, we did not observe a significant trialwise correlation of the SNR metric with microsaccade count in either direction. This might imply that either the induced gamma power adds both to signal and to the noise term and hence does not affect the SNR measure, or that the effect of microsaccadic gamma on SNR is too weak (compared to other activity) to manifest in significant changes of SNR. Furthermore previous research has shown that the effect of spike potentials and eyelid-induced signal changes is strongest on frontal channels (Plöchl et al., 2012), while the topography contrasts of our ASSR measure revealed that effects of the fixation task were only significant in occipital channels (see Fig. 4e). Recent work using intracranial EEG recordings over occipital cortex areas presented evidence for a characteristic gamma decrease/increase pattern related to eye blinks and larger saccadic eye movements (Kern et al., 2021). The pattern was discussed as reflecting a potential neuronal mechanism for active visual suppression in the visual cortex in order to achieve stable perception. As gamma effects with these sources would fit the observed topography in our data it appears possible that this process could affect the 40 Hz SNR measure. However, the effects were only observed in invasive recordings that offer much higher signal quality than scalp EEG, and were reported for gamma band above 50 Hz. It is unclear whether the described effects would also be present around 40 Hz and whether they would even be detectable in scalp EEG. Furthermore in our data, ICA based cleaning of the EEG removed the effect of fixation task on ASSR; the applied classifier was trained on a large set of ICA components labelled by experts who would be unlikely to label an ICA component located over the visual cortex as noise. It therefore seems likely that the ASSR metric is indeed sensitive to a broader set of artifactual signal distortions, even if they do not result in significantly different SNR values for specific channels. Further research would be necessary to validate this interpretation, and also to investigate a potential interaction of 40 Hz ASSR and broadband gamma activity related to eye movements and early visual processing.

Lastly, the question remains whether the brain’s actual ASSR (not its measured SNR at scalp level) might be systematically affected by factors not controlled for in the present study. Directed attention towards the auditory stream is controversially discussed to influence the strength of ASSR: while early work found no overall effect of selective attention on ASSR (attending the stimulus vs. reading a book; Linden et al., 1987), more recent work did (e.g. for attention shifts between two auditory stimuli [Skosnik et al. 2007] or for attention shift from the visual to the auditory domain or vice versa [Saupe et al. 2009]). If this were true it constitutes a potential confound for the proxy measure only if attention to the auditory stream should be systematically linked to the investigated primary task. In the present study, we took measures to avoid such a link and didn’t find significant evidence for any attention-related confound. However, the topography contrasts in Fig. 4e show hints of potential sub-threshold effects on the temporal dipoles that might mark sources of the ASSR, most clearly in the content condition (increased polarity of dipoles in dance vs. landscape scenes). This could be interpreted as consistent with attentional suppression or enhancement, if attention were to be shifted away from the auditory stimulus onto the visual stimulus for landscape scenes or toward the auditory stimulus for dance scenes, leading to suppression of the ASSR. In our behavioral data, however, we see no clear evidence that such an attentional shift occurred: aesthetic appeal was rated significantly higher in landscape scenes than in dance scenes, which would be expected to cause a shift of attention away from the auditory to the visual modality. Yet on a trial-by-trial basis aesthetic appeal correlated negatively with SNR (if at all), which contradicts this interpretation. In addition, the differences between conditions observed at these temporal electrodes were quite small, failing to reach significance; it would therefore be prudent to avoid over-interpreting the observed pattern.

### 4.7 Conclusion

As cognitive neuroscience proceeds to address more complex and integrative mental processes such as aesthetic experiences, the field will need to apply increasingly naturalistic experimental settings and stimuli. Using a novel measure of EEG signal quality, we find that one of the most common experimental constraints in visual cognitive neuroscience — the use of static visual stimuli — can potentially be relaxed without significant signal loss, and that doing so can have a number of positive effects on behavior and physiology. These findings are encouraging for future work using EEG in more naturalistic paradigms. We hope that our work will help researchers find the right balance between ecological validity on the one hand and the need for high signal quality and interpretability of findings on the other.

## 5 Additional information

### 5.1 Data and code availability

The experimental design and hypotheses reported here were preregistered (https://osf.io/bkep4). The manuscript was uploaded to a preprint server (https://www.biorxiv.org/content/10.1101/2021.09.18.460905). The dataset for statistical analysis, as well as R and python scripts replicating the results, and python scripts used in data collection were made available in a public online repository (https://osf.io/8zctp). The stimulus set contains copyright protected material and can not be publicly shared. Raw EEG data can not be publicly shared due to ethics regulations at our institute. The code created for computing ASSR SNR was also made publicly available as part of a tutorial in the MNE-Python documentation; https://mne.tools/stable/auto_tutorials/time-freq/50_ssvep.html.

## 5.2 Acknowledgements

The authors thank Ayse Ilkay Isik for the original stimulus set, Sarah Koldehoff for cutting the final video stimuli, Elke Lange and Lauren Fink for their valuable comments and feedback on the Eyetracking analysis, and to Sarah Koldehoff, and Johannes Messerschmidt for assistance with the data collection. The work was supported by the Max Planck Society.

## 5.3 Author statement (CRediT)

DW: Conceptualization, Methodology, Investigation, Formal Analysis, Software, Data curation, Project administration, Visualization, Writing - Original Draft, Writing - Review & Editing EV: Conceptualization, Writing - Original Draft, Writing - Review & Editing, Resources, Supervision, Funding acquisition

## Supplementary Material

### Participant demographics

See Tabs. 2 and 3 for full sample demographics.

**Table 2:**
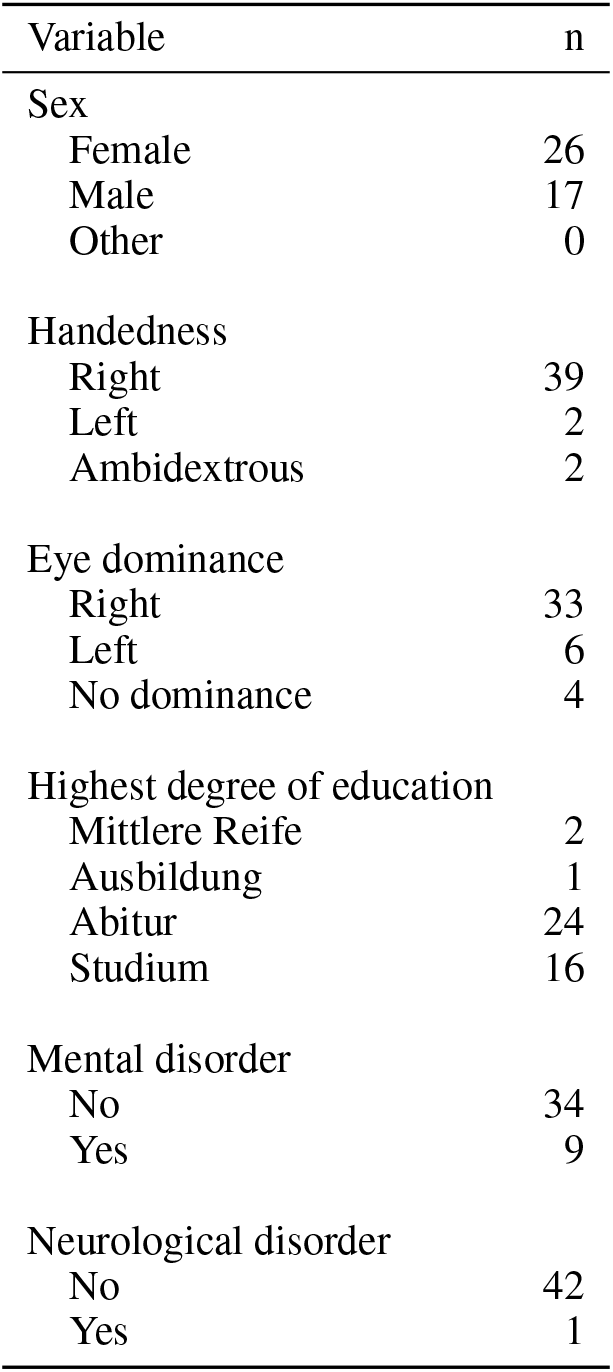
Categorical demographic factors

**Table 3:** Continuous demographic factors

| Variable | n | M | SD | min | max | 25% | 50% | 75% |
| --- | --- | --- | --- | --- | --- | --- | --- | --- |
| Age | 43 | 27.1 | 7.06 | 19 | 52 | 22.5 | 25 | 30 |
| Years of education | 43 | 17.8 | 3.55 | 9 | 25 | 15 | 18 | 20 |
| Caffeine intake [mg/kg] | 43 | 0.84 | 1.041 | 0.00 | 4.28 | 0.00 | 0.52 | 1.43 |
| BFI (range 3-15) |  |  |  |  |  |  |  |  |
| extraversion | 43 | 9.30 | 1.897 | 5 | 13 | 8 | 9 | 10.5 |
| open mindedness | 43 | 12.14 | 1.684 | 8 | 15 | 11 | 13 | 13 |
| agreeableness | 43 | 11.28 | 1.894 | 7 | 15 | 10 | 12 | 12.5 |
| conscientiousness | 43 | 10.65 | 2.516 | 5 | 15 | 9 | 11 | 13 |
| negative emotionality | 43 | 8.23 | 2.983 | 3 | 15 | 6 | 8 | 10 |
| PANAS (range 6-30) |  |  |  |  |  |  |  |  |
| positive | 43 | 17.60 | 4.588 | 9 | 25 | 14 | 18 | 21 |
| negative | 43 | 7.47 | 2.364 | 6 | 15 | 6 | 6 | 7.5 |
| SHAPS (range 0-14) | 43 | 1.02 | 1.318 | 0 | 5 | 0 | 1 | 2 |
| BPS (range 8-56) | 43 | 19.07 | 8.213 | 8 | 49 | 15.5 | 17 | 21.5 |

### Log transformation of ASSR SNR

SNR values were log transformed to shift them from a skewed gamma to a more gaussian distribution (see Fig. 5). We applied the natural logarithm to average SNR values over all EEG channels for each participant and each trial.

**Figure 5:**
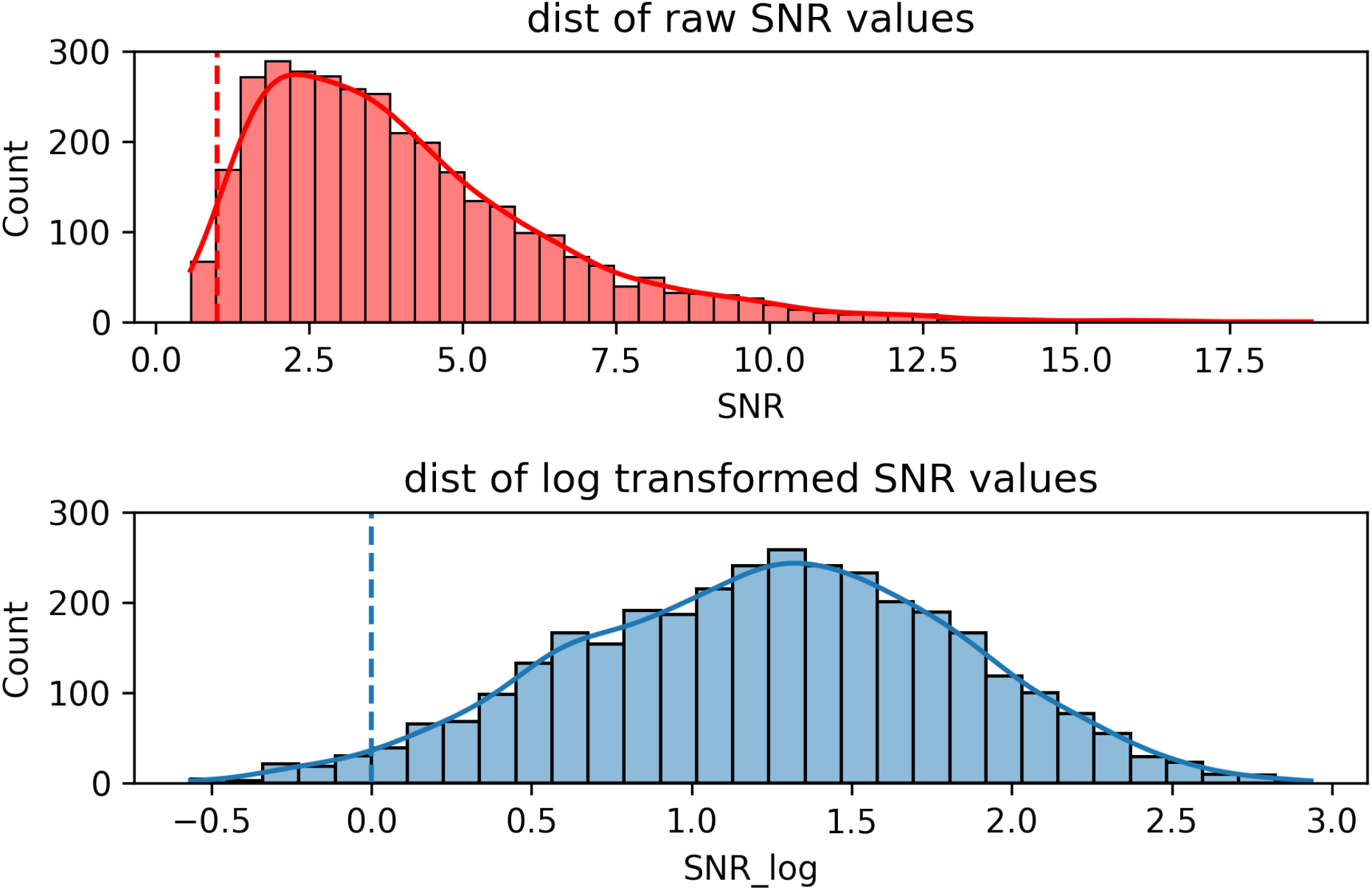
Distribution of raw and log transformed SNR values across all participants (N=43)

### Full ANOVA tables

Results of all repeated measures ANOVA models in the study are compiled in Tab. 4.

**Table 4:**
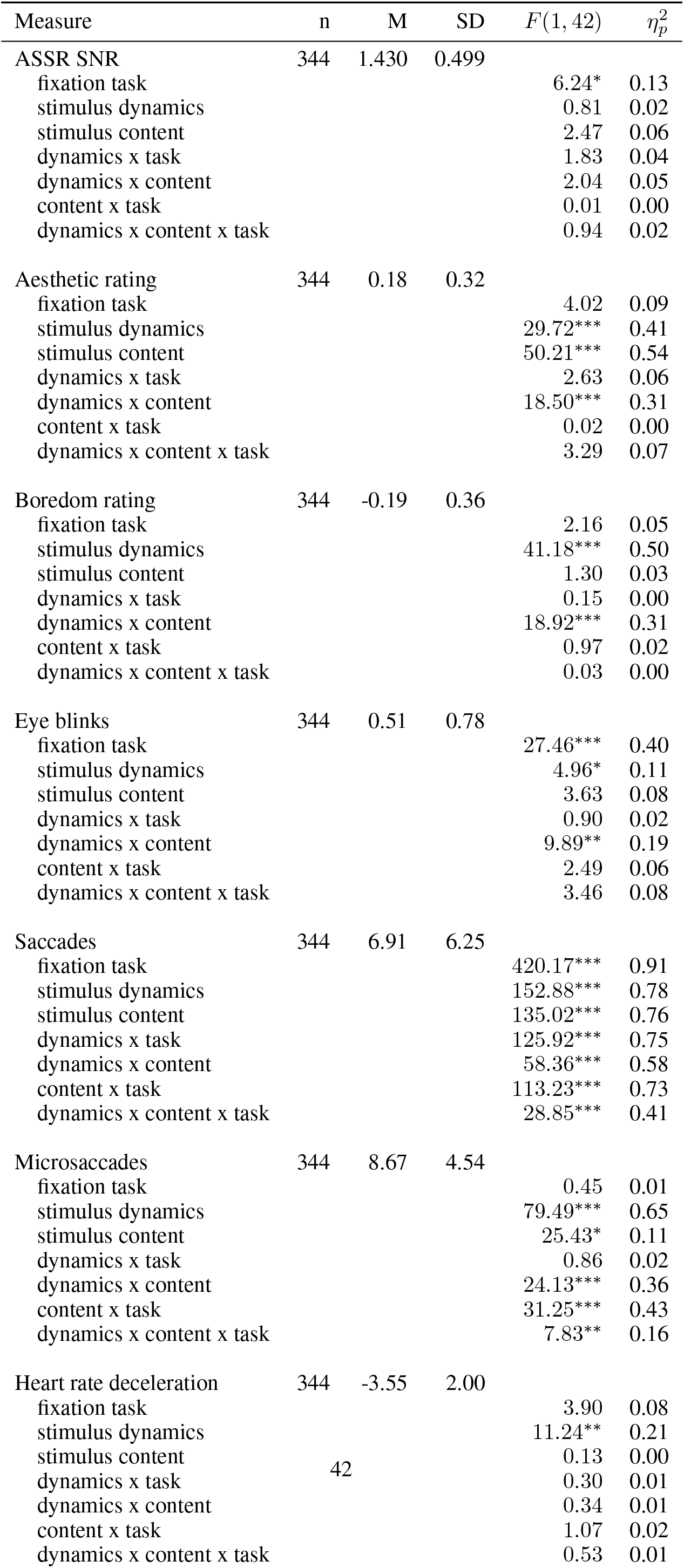
Full ANOVA results

### Trial wise correlation of all dependent measures

See Tab. 5

**Table 5:** Full correlation table for trialwise measures

| Measure | n | M | SD | 1 | 2 | 3 | 4 | 5 | 6 |
| --- | --- | --- | --- | --- | --- | --- | --- | --- | --- |
| ASSR SNR | 3349 | 1.43 | 0.64 | — |  |  |  |  |  |
| Ratings |  |  |  |  |  |  |  |  |  |
| aesthetic | 3349 | .19 | .53 | -.046 | — |  |  |  |  |
| boredom | 3349 | -.19 | .58 | .038 | -.580*** | — |  |  |  |
| Eye movements |  |  |  |  |  |  |  |  |  |
| eye blinks | 3189 | 0.54 | 1.06 | -.053* | -.059* | .078*** | — |  |  |
| saccades | 3349 | 7.03 | 6.91 | -.041 | .018 | .029 | — | — |  |
| microsaccades | 3349 | 8.64 | 6.01 | .000 | -.086*** | .126*** | — | — | — |
| Heart rate change (bpm) | 3336 | -3.57 | 4.48 | -.036 | .005 | -.010 | .049 | .059* | -.000 |

*Note:* $r$ values computed using repeated measures correlation (Bakdash & Marusich, 2017). \* $p < .05$ , \*\*\* $p < .001$ . $p$ values corrected using Holm's method (Holm, 1979). $n$ trials for eye blinks is lower because two participants exhibited zero blinks over all trials, which prevented the model to converge; $n$ trials for HR deceleration is lower because outliers were rejected. Correlation between the different eye measures was not investigated.

### Frequency dependent correlation of eye tracking signal and EEG

As it is unclear whether our proxy measure for signal quality, SNR of the 40 Hz ASSR, might be confined to detect noise only of a specific frequency characteristic (e.g. close to the low gamma range) we wanted to investigate another frequency dependant measures for noise detection. Given the well known physiological effect of eye movements on EEG, we want to test whether the time-varying band power in a given frequency band of the EEG signal is correlated with band power of the EOG in the same frequency band. This might hint at an induction of artifactual eye movement related signal into the EEG, thereby potentially confounding frequency based analysis. Here we correlated global field power (GFP) of EEG and eye tracking data in the commonly used frequency bands delta (1-3 Hz), theta (4-7 Hz), alpha (8-12 Hz), beta (13-25 Hz), low gamma (30-45 Hz), and high gamma (45 – 120 Hz).

#### Methods

We largely followed the analysis as described in (Engemann & Appelhoff, n.d.), which uses an adapted version of the method described in (Hari, 1997).

As we did not record EOG, X and Y coordinates of the fixation time series from the calibrated eye tracking data were taken as the raw data to indicate eyeball movements. These two signals can be expected to carry similar information, as changes in X and Y fixation location on the screen are caused by horizontal and vertical eye movements and directly reflect eyeball rotation. Missing values (e.g. caused by blinks) were exchanged with zeros. EEG data were down-sampled to 500 Hz, the sampling frequency of the eye tracker.

First EEG and ET data were bandpass filtered according to the respective frequency band (FIR, zero-phase, 1 Hz transition band, filter length automatically chosen based on the size of the transition regions). Then a Hilbert transform was applied. Next, the evoked response was subtracted from every single trial of the EEG to reveal oscillatory activity (but not from the eye tracking signal). The signals were rectified per channel by taking the magnitude of the Hilbert transform. Then the GFP was computed using the sum of squares, across all channels (EEG) and across left and right eye channels of binocular recordings respectively. The procedure was repeated for each frequency band of interest and the correlation between EEG GFP on the one hand and GFP of vertical and horizontal component of the eye movements on the other hand was computed for each participant. Repeated measures correlation (Bakdash & Marusich, 2017) was used to account for the trial structure of the data (using the trial number as grouping variable). This procedure was conducted with all trials from all conditions. In a last step, average correlation coefficients across participants were calculated for all frequency bands. Significance statements are difficult, as the multiple comparison problem cannot be addressed in a straightforward manner. We decided for the following approach: as individual correlation coefficients on the participant level have corresponding p-values, these were corrected for multiple comparison using Holm’s method (Holm, 1979), and the average Holm-corrected p-value as well as the percentage of significant data points is reported for each frequency band. However, we think that the resulting significance values should not be over-interpreted.

#### Results

Tab. 6 summarizes the results of the correlation analysis. We see that there is indeed a substantial correlation between oscillatory dynamics in EEG and Eye movements in certain frequency bands. Especially in the lower frequencies (Delta, Theta) the two signals correlate very strongly, while the correlation falls off steeply in the medium and high frequency bands starting with the alpha-band around 8 Hz. Qualitatively, horizontal eye movements seem to be slightly stronger linked to EEG band power dynamics.

**Table 6:** Correlation between global field power of EEG and eye tracking (ET) signal in different frequency bands

| Frequency band | EEG x horizontal ET | EEG x vertical ET |
| --- | --- | --- |
| Delta (1-4 Hz) | .68*** | .61* |
| Theta (4-7 Hz) | .51*** | .47* |
| Alpha (8-12 Hz) | .10* | .10* |
| Beta (13-25 Hz) | .03 | .03 |
| Gamma low (30-45 Hz) | .01 | .01 |
| Gamma high (46-120 Hz) | .01 | .01 |

*Note:* $r$ values computed using repeated measures correlation (Bakdash & Marusich, 2017) across trials for each participants; depicted values are grand averages across all participants. $p$ -values for each participant corrected using Holm's method (Holm, 1979) and averaged across participants. \* $p < .05$ , \*\*\* $p < .001$ .

Average Holm-corrected p-values across participants are reported in Tab. 6. Most of the correlation coefficients (per participant) between eye movements and EEG were significant after multiple-comparison correction (*p* < .05): 84.9 % for horizontal eye movements and 82.2 % for vertical eye movements.

#### Discussion

In general, the analysis shows that in our data oscillatory dynamics of EEG and eye movements can be linked, depending on the frequency band. This is in line with an induction of artifactual signal into the EEG, which was already known from the literature. Especially in the lower frequencies (delta and theta) the two signals correlate very strongly, which might reflect the large signal offsets induced in the EEG by rotation of the eyes’ dipoles (this offset was shown to increased linearly with the size of the eye movement; Plöchl et al., 2012). This finding is of particular relevance for research interested in oscillatory dynamics in these low frequencies: for such studies, removing the fixation task and refraining from rejecting trials with eye movements might not be an option, as it bears the risk of misinterpreting artifacts induced by eye movements as neuronal effects. We want to note though, that additional cleaning of the EEG using ICA or similar approaches might well remove or reduce the correlation of the two signals; we did not explicitly test for this, though.

High frequent EEG artifacts in the gamma band, induced by e.g. saccadic spike potentials or microsaccades (see Plöchl et al., 2012; Yuval-Greenberg et al., 2008), were not reflected by a correlation between the two signals, despite the fact that there must have been eye movements during each single trial, regardless of the condition (see Sec. 3 eye tracking). Apparently, the GFP correlation method is insensitive to these high frequent eye movement artifacts. This might be due to the fact that the raw eye trace, as opposed to EOG proper, does not contain thes signal components; unfortunately, our dataset does not allow to test for this. Our ASSR SNR measure, on the other hand, did correlate with blinks and larger saccades, but not with microsaccades. It seems possible that ASSR SNR mainly reacts to these high frequent noise components, as they are in the same frequency range as the signal (40 Hz), even though more broadband. If this where true, it would question the usability of the proxy metric. However, previous research has shown that the effect of spike potentials and eyelid-induced signal changes is strongest on frontal channels (Plöchl et al., 2012), while the topography contrasts of our ASSR measure revealed that effects of the fixation task were only significant in occipital channels (see Fig. 4e). We thus do believe that the ASSR metric is sensitive to a broader set of artifactual signal distortions.

The small correlation of the signals in the alpha and beta band might be less of a concern for future research. There might be some amount of eye induced signal reflected in this frequency range, but it seems unlikely that this might flaw an entire study, especially if approaches like trial averaging or signal cleaning are involved.

On a sidenote, previous work indicated that artifacts caused by vertical eye movements would have a higher influence on the EEG (Plöchl et al., 2012). In this analysis however, the correlation of EEG bandpower with the horizontal component of the eye movements was stronger than with the vertical component.

## Footnotes

1 Note that algorithm based data cleaning can leave residuals in the data (see Dimigen, 2020; Robbins et al., 2020, for ICA), which can be especially problematic if subsequent analyses are based on machine learning approaches (Quax et al., 2019; Thielen et al., 2019).

## References

Appelhoff, S., Sanderson, M., Brooks, T., van Vliet, M., Quentin, R., Holdgraf, C., Chaumon, M., Mikulan, E., Tavabi, K., Höchenberger, R., Welke, D., Brunner, C., Rockhill, A., Larson, E., Gramfort, A., & Jas, M. (2019). MNE-BIDS: Organizing electrophysiological data into the BIDS format and facilitating their analysis. Journal of Open Source Software, 4(44), 1896. https://doi.org/10.21105/joss.01896

Armstrong, T., & Detweiler-Bedell, B. (2008). Beauty as an emotion: The exhilarating prospect of mastering a challenging world. Review of General Psychology, 12(4), 305–329. https://doi.org/10.1037/a0012558

Auerbach-Asch, C. R., Bein, O., & Deouell, L. Y. (2020). Face Selective Neural Activity: Comparisons Between Fixed and Free Viewing. Brain Topography, 33(3), 336–354.https://doi.org/10.1007/s10548-020-00764-7

Ayrolles, A., Brun, F., Chen, P., Djalovski, A., Beauxis, Y., Delorme, R., Bourgeron, T., Dikker, S., & Dumas, G. (2021). HyPyP: A Hyperscanning Python Pipeline for inter-brain connectivity analysis. Social Cognitive and Affective Neuroscience, 16(1-2), 72–83. https://doi.org/10.1093/scan/nsaa141

Bakdash, J. Z., & Marusich, L. R. (2017). Repeated Measures Correlation. Frontiers in Psychology, 8. https://doi.org/10.3389/fpsyg.2017.00456

Ball, T., Kern, M., Mutschler, I., Aertsen, A., & Schulze-Bonhage, A. (2009). Signal quality of simultaneously recorded invasive and non-invasive EEG. NeuroImage, 46(3), 708–716.https://doi.org/10.1016/j.neuroimage.2009.02.028

Bergamasco, B., Benna, P., Covacich, A. M., Gilli, M., & Rossi, G. (1976). Behaviour Of CNV During Exposure To Urban Traffic Noise. Acta Oto-Laryngologica, 81(sup339), 27–29. https://doi.org/10.3109/00016487609124919

Bigdely-Shamlo, N., Mullen, T., Kothe, C., Su, K.-M., & Robbins, K. A. (2015). The PREP pipeline: Standardized preprocessing for large-scale EEG analysis. Frontiers in Neuroinformatics, 9. https://doi.org/10.3389/fninf.2015.00016

Bohórquez, J., & Özdamar, Ö. (2008). Generation of the 40-Hz auditory steady-state response (ASSR) explained using convolution. Clinical Neurophysiology, 119(11), 2598–2607.https://doi.org/10.1016/j.clinph.2008.08.002

Braboszcz, C., & Delorme, A. (2011). Lost in thoughts: Neural markers of low alertness during mind wandering. NeuroImage, 54(4), 3040–3047.https://doi.org/10.1016/j.neuroimage.2010.10.008

Brieber, D., Nadal, M., & Leder, H. (2015). In the white cube: Museum context enhances the valuation and memory of art. Acta Psychologica, 154, 36–42. https://doi.org/10.1016/j.actpsy.2014.11.004

Brieber, D., Nadal, M., Leder, H., & Rosenberg, R. (2014). Art in Time and Space: Context Modulates the Relation between Art Experience and Viewing Time (L. M. Martinez, Ed.). PLoS ONE, 9(6), e99019. https://doi.org/10.1371/journal.pone.0099019

Brielmann, A. A., & Pelli, D. G. (2017). Beauty requires thought. Current Biology, 27(10), 1506–1513.e3. https://doi.org/10.1016/j.cub.2017.04.018

Bühler, E., Lachenmeier, D. W., & Winkler, G. (2014). Development of a tool to assess caffeine intake among teenagers and young adults. Ernahrungs Umschau, 61(4), 58–63.https://doi.org/10.4455/eu.2014.011

Chatterjee, A., & Vartanian, O. (2014). Neuroaesthetics. Trends in Cognitive Sciences, 18(7), 370–375. https://doi.org/10.1016/j.tics.2014.03.003

Coles, M. G. H. (1972). Cardiac and respiratory activity during visual search. Journal of Experimental Psychology, 96(2), 371–379. https://doi.org/10.1037/h0033603

Coll, M.-P., Hobson, H., Bird, G., & Murphy, J. (2021). Systematic review and meta-analysis of the relationship between the heartbeat-evoked potential and interoception. Neuroscience & Biobehavioral Reviews, 122, 190–200. https://doi.org/10.1016/j.neubiorev.2020.12.012

Committee. (1958). Report of the committee on methods of clinical examination in electroencephalography. Electroencephalography and Clinical Neurophysiology, 10(2), 370–375.https://doi.org/10.1016/0013-4694(58)90053-1

Craddock, M., Martinovic, J., & Müller, M. M. (2016). Accounting for microsaccadic artifacts in the EEG using independent component analysis and beamforming: ICA, beamforming, and artifactual gamma. Psychophysiology, 53(4), 553–565.https://doi.org/10.1111/psyp.12593

Debener, S., Minow, F., Emkes, R., Gandras, K., & de Vos, M. (2012). How about taking a low-cost, small, and wireless EEG for a walk?: EEG to go. Psychophysiology, 49(11), 1617–1621.https://doi.org/10.1111/j.1469-8986.2012.01471.x

Delorme, A., & Makeig, S. (2004). EEGLAB: An open source toolbox for analysis of single-trial EEG dynamics including independent component analysis. Journal of Neuroscience Methods, 134(1), 9–21. https://doi.org/10.1016/j.jneumeth.2003.10.009

Dias, J. C., Sajda, P., Dmochowski, J. P., & Parra, L. C. (2013). EEG precursors of detected and missed targets during free-viewing search. Journal of Vision, 13(13), 13–13.https://doi.org/10.1167/13.13.13

Dikker, S., Michalareas, G., Oostrik, M., Serafimaki, A., Kahraman, H. M., Struiksma, M. E., & Poeppel, D. (2021). Crowdsourcing neuroscience: Inter-brain coupling during face-to-face interactions outside the laboratory. NeuroImage, 227, 117436. https://doi.org/10.1016/j.neuroimage.2020.117436

Dikker, S., Wan, L., Davidesco, I., Kaggen, L., Oostrik, M., McClintock, J., Rowland, J., Michalareas, G., Van Bavel, J. J., Ding, M., & Poeppel, D. (2017). Brain-to-Brain Synchrony Tracks Real-World Dynamic Group Interactions in the Classroom. Current Biology, 27(9), 1375–1380. https://doi.org/10.1016/j.cub.2017.04.002

Dimigen, O. (2020). Optimizing the ICA-based removal of ocular EEG artifacts from free viewing experiments. NeuroImage, 207, 116–117. https://doi.org/10.1016/j.neuroimage.2019.116117

Dimigen, O., Sommer, W., Hohlfeld, A., Jacobs, A. M., & Kliegl, R. (2011). Coregistration of eye movements and EEG in natural reading: Analyses and review. Journal of Experimental Psychology: General, 140(4), 552–572. https://doi.org/10.1037/a0023885

Dmochowski, J. P., Bezdek, M. A., Abelson, B. P., Johnson, J. S., Schumacher, E. H., & Parra, L. C. (2014). Audience preferences are predicted by temporal reliability of neural processing. Nature Communications, 5(1). https://doi.org/10.1038/ncomms5567

Dmochowski, J. P., Sajda, P., Dias, J., & Parra, L. C. (2012). Correlated Components of Ongoing EEG Point to Emotionally Laden Attention – A Possible Marker of Engagement? Frontiers in Human Neuroscience, 6. https://doi.org/10.3389/fnhum.2012.00112

Eckstein, M. K., Guerra-Carrillo, B., Miller Singley, A. T., & Bunge, S. A. (2017). Beyond eye gaze: What else can eyetracking reveal about cognition and cognitive development? Developmental Cognitive Neuroscience, 25, 69–91. https://doi.org/10.1016/j.dcn.2016.11.001

Ehinger, B. V., & Dimigen, O. (2019). Unfold: An integrated toolbox for overlap correction, non-linear modeling, and regression-based EEG analysis. PeerJ, 7, e7838. https://doi.org/10.7717/peerj.7838

Engbert, R., & Mergenthaler, K. (2006). Microsaccades are triggered by low retinal image slip. Proceedings of the National Academy of Sciences, 103(18), 7192–7197.https://doi.org/10.1073/pnas.0509557103

Engemann, D. A., & Appelhoff, S. (n.d.). Explore event-related dynamics for specific frequency bands. Retrieved January 20, 2022, from https://mne.tools/stable/auto_examples/time_frequency/time_frequency_global_field_power.html

Faul, F., Erdfelder, E., Lang, A.-G., & Buchner, A. (2007). G*Power 3: A flexible statistical power analysis program for the social, behavioral, and biomedical sciences. Behavior Research Methods, 39(2), 175–191.https://doi.org/10.3758/BF03193146

Fink, L. K., Lange, E. B., & Groner, R. (2019). The application of eye-tracking in music research. Journal of Eye Movement Research, 11(2), 1. https://doi.org/10.16910/jemr.11.2.1

Fox, K. C., Spreng, R. N., Ellamil, M., Andrews-Hanna, J. R., & Christoff, K. (2015). The wandering brain: Metaanalysis of functional neuroimaging studies of mind-wandering and related spontaneous thought processes. NeuroImage, 111, 611–621. https://doi.org/10.1016/j.neuroimage.2015.02.039

Galambos, R., Makeig, S., & Talmachoff, P. J. (1981). A 40-Hz auditory potential recorded from the human scalp. Proceedings of the National Academy of Sciences, 78(4), 2643–2647.https://doi.org/10.1073/pnas.78.4.2643

Goldstein, R. B., Woods, R. L., & Peli, E. (2007). Where people look when watching movies: Do all viewers look at the same place? Computers in Biology and Medicine, 37(7), 957–964. https://doi.org/10.1016/j.compbiomed.2006.08.018

Gramann, K., Ferris, D. P., Gwin, J., & Makeig, S. (2014). Imaging natural cognition in action. International Journal of Psychophysiology, 91(1), 22–29.https://doi.org/10.1016/j.ijpsycho.2013.09.003

Gramann, K., Hohlefeld, F. U., Gehrke, L., & Klug, M. (2021). Human cortical dynamics during full-body heading changes. Scientific Reports, 11(1), 18186. https://doi.org/10.1038/s41598-021-97749-8

Gramfort, A. (2013). MEG and EEG data analysis with MNE-Python. Frontiers in Neuroscience, 7. https://doi.org/10.3389/fnins.2013.00267

Greeno, J. G. (1994). Gibson’s affordances. Psychological Review, 101(2), 336–342.https://doi.org/10.1037/0033-295X.101.2.336

Gwin, J. T., Gramann, K., Makeig, S., & Ferris, D. P. (2011). Electrocortical activity is coupled to gait cycle phase during treadmill walking. NeuroImage, 54(2), 1289–1296.https://doi.org/10.1016/j.neuroimage.2010.08.066

Hari, R. (1997). Human cortical oscillations: A neuromagnetic view through the skull. Trends in Neurosciences, 20(1), 44–49.https://doi.org/10.1016/S0166-2236(96)10065-5

Hasson, U. (2004). Intersubject Synchronization of Cortical Activity During Natural Vision. Science, 303(5664), 1634–1640.https://doi.org/10.1126/science.1089506

Hasson, U., & Frith, C. D. (2016). Mirroring and beyond: Coupled dynamics as a generalized framework for modelling social interactions. Philosophical Transactions of the Royal Society B: Biological Sciences, 371(1693), 20150366. https://doi.org/10.1098/rstb.2015.0366

Hasson, U., & Honey, C. J. (2012). Future trends in Neuroimaging: Neural processes as expressed within real-life contexts. NeuroImage, 62(2), 1272–1278.https://doi.org/10.1016/j.neuroimage.2012.02.004

Heimann, K., Umiltà, M. A., Guerra, M., & Gallese, V. (2014). Moving Mirrors: A High-density EEG Study Investigating the Effect of Camera Movements on Motor Cortex Activation during Action Observation. Journal of Cognitive Neuroscience, 26(9), 2087–2101.https://doi.org/10.1162/jocn_a_00602

Holm, S. (1979). A Simple Sequentially Rejective Multiple Test Procedure. Scandinavian Journal of Statistics, 6(2), 65–70.

Isik, A. I., & Vessel, E. A. (2019). Continuous ratings of movie watching reveal idiosyncratic dynamics of aesthetic enjoyment (R. Ferrer, Ed.). PLOS ONE, 14(10), e0223896. https://doi.org/10.1371/journal.pone.0223896

Isik, A. I., & Vessel, E. A. (2021). From visual perception to aesthetic appeal: Brain responses to aesthetically appealing natural landscape movies. Frontiers in Human Neuroscience.

Iwasaki, M., Kellinghaus, C., Alexopoulos, A. V., Burgess, R. C., Kumar, A. N., Han, Y. H., Lüders, H. O., & Leigh, R. J. (2005). Effects of eyelid closure, blinks, and eye movements on the electroencephalogram. Clinical Neurophysiology, 116(4), 878–885. https://doi.org/10.1016/j.clinph.2004.11.001

Johnston, P., Molyneux, R., & Young, A. W. (2015). The N170 observed ‘in the wild’: Robust event-related potentials to faces in cluttered dynamic visual scenes. Social Cognitive and Affective Neuroscience, 10(7), 938–944. https://doi.org/10.1093/scan/nsu136

Jongkees, B. J., & Colzato, L. S. (2016). Spontaneous eye blink rate as predictor of dopamine-related cognitive function—A review. Neuroscience & Biobehavioral Reviews, 71, 58–82. https://doi.org/10.1016/j.neubiorev.2016.08.020

Kam, J. W. Y., Dao, E., Farley, J., Fitzpatrick, K., Smallwood, J., Schooler, J. W., & Handy, T. C. (2011). Slow Fluctuations in Attentional Control of Sensory Cortex. Journal of Cognitive Neuroscience, 23(2), 460–470.https://doi.org/10.1162/jocn.2010.21443

Kam, J. W. Y., Irving, Z. C., Mills, C., Patel, S., Gopnik, A., & Knight, R. T. (2021). Distinct electrophysiological signatures of task-unrelated and dynamic thoughts. Proceedings of the National Academy of Sciences, 118(4), e2011796118. https://doi.org/10.1073/pnas.2011796118

Kamienkowski, J. E., Ison, M. J., Quiroga, R. Q., & Sigman, M. (2012). Fixation-related potentials in visual search: A combined EEG and eye tracking study. Journal of Vision, 12(7), 4–4. https://doi.org/10.1167/12.7.4

Kaunitz, L. N., Kamienkowski, J. E., Varatharajah, A., Sigman, M., Quiroga, R. Q., & Ison, M. J. (2014). Looking for a face in the crowd: Fixation-related potentials in an eye-movement visual search task. NeuroImage, 89, 297–305. https://doi.org/10.1016/j.neuroimage.2013.12.006

Keitel, C., Maess, B., Schröger, E., & Müller, M. M. (2013). Early visual and auditory processing rely on modality-specific attentional resources. NeuroImage, 70, 240–249. https://doi.org/10.1016/j.neuroimage.2012.12.046

Kern, M., Schulze-Bonhage, A., & Ball, T. (2021). Blink-and saccade-related suppression effects in early visual areas of the human brain: Intracranial EEG investigations during natural viewing conditions. NeuroImage, 230, 117788. https://doi.org/10.1016/j.neuroimage.2021.117788

Kovach, C. K., Tsuchiya, N., Kawasaki, H., Oya, H., Howard, M. A., & Adolphs, R. (2011). Manifestation of ocular-muscle EMG contamination in human intracranial recordings. NeuroImage, 54(1), 213–233. https://doi.org/10.1016/j.neuroimage.2010.08.002

Kriegeskorte, N. (2008). Representational similarity analysis – connecting the branches of systems neuroscience. Frontiers in Systems Neuroscience. https://doi.org/10.3389/neuro.06.004.2008

Lacourse, K., Yetton, B., Mednick, S., & Warby, S. C. (2020). Massive online data annotation, crowdsourcing to generate high quality sleep spindle annotations from EEG data. Scientific Data, 7(1), 190. https://doi.org/10.1038/s41597-020-0533-4

Lakens, D. (2013). Calculating and reporting effect sizes to facilitate cumulative science: A practical primer for t-tests and ANOVAs. Frontiers in Psychology, 4. https://doi.org/10.3389/fpsyg.2013.00863

Linden, R. D., Picton, T. W., Hamel, G., & Campbell, K. B. (1987). Human auditory steady-state evoked potentials during selective attention. Electroencephalography and Clinical Neurophysiology, 66(2), 145–159.https://doi.org/10.1016/0013-4694(87)90184-2

Liu, C., Herrup, K., Goto, S., & Shi, B. (2020). Viewing garden scenes: Interaction between gaze behavior and physiological responses. Journal of Eye Movement Research, 13(1). https://doi.org/10.16910/jemr.13.1.6

Lopez-Persem, A., Bastin, J., Petton, M., Abitbol, R., Lehongre, K., Adam, C., Navarro, V., Rheims, S., Kahane, P., Domenech, P., & Pessiglione, M. (2020). Four core properties of the human brain valuation system demonstrated in intracranial signals. Nature Neuroscience. https://doi.org/10.1038/s41593-020-0615-9

Lu, Z., & Ku, Y. (2020). NeuroRA: A Python Toolbox of Representational Analysis From Multi-Modal Neural Data. Frontiers in Neuroinformatics, 14, 563669. https://doi.org/10.3389/fninf.2020.563669

Luck, S. J. (2014). An introduction to the event-related potential technique (Second edition) [OCLC: 896211347]. The MIT Press.

Luck, S. J., Stewart, A. X., Simmons, A. M., & Rhemtulla, M. (2021). Standardized measurement error: A universal metric of data quality for averaged event-related potentials. Psychophysiology. https://doi.org/10.1111/psyp.13793

Lustenberger, C., Patel, Y. A., Alagapan, S., Page, J. M., Price, B., Boyle, M. R., & Fröhlich, F. (2018). High-density EEG characterization of brain responses to auditory rhythmic stimuli during wakefulness and NREM sleep. NeuroImage, 169, 57–68. https://doi.org/10.1016/j.neuroimage.2017.12.007

Makeig, S., Gramann, K., Jung, T.-P., Sejnowski, T. J., & Poizner, H. (2009). Linking brain, mind and behavior. International Journal of Psychophysiology, 73(2), 95–100.https://doi.org/10.1016/j.ijpsycho.2008.11.008

Maris, E., & Oostenveld, R. (2007). Nonparametric statistical testing of EEG-and MEG-data. Journal of Neuroscience Methods, 164(1), 177–190. https://doi.org/10.1016/j.jneumeth.2007.03.024

McVay, J. C., & Kane, M. J. (2012). Drifting from slow to “d’oh!”: Working memory capacity and mind wandering predict extreme reaction times and executive control errors. Journal of Experimental Psychology: Learning, Memory, and Cognition, 38(3), 525–549.https://doi.org/10.1037/a0025896

Meigen, T., & Bach, M. (1999). On the statistical significance of electrophysiological steady-state responses. Documenta Ophthalmologica, 98(3), 207–232.https://doi.org/10.1023/A:1002097208337

Menninghaus, W., Wagner, V., Hanich, J., Wassiliwizky, E., Kuehnast, M., & Jacobsen, T. (2015). Towards a psychological construct of being moved (H. Nusbaum, Ed.). PLOS ONE, 10(6), e0128451. https://doi.org/10.1371/journal.pone.0128451

Merrifield, C., & Danckert, J. (2014). Characterizing the psychophysiological signature of boredom. Experimental Brain Research, 232(2), 481–491. https://doi.org/10.1007/s00221-013-3755-2

Mitrovic, A., Hegelmaier, L. M., Leder, H., & Pelowski, M. (2020). Does beauty capture the eye, even if it’s not (overtly) adaptive? A comparative eye-tracking study of spontaneous attention and visual preference with VAST abstract art. Acta Psychologica, 209, 103133. https://doi.org/10.1016/j.actpsy.2020.103133

Nikolaev, A. R., Meghanathan, R. N., & van Leeuwen, C. (2016). Combining EEG and eye movement recording in free viewing: Pitfalls and possibilities. Brain and Cognition, 107, 55–83. https://doi.org/10.1016/j.bandc.2016.06.004

Norcia, A. M., Appelbaum, L. G., Ales, J. M., Cottereau, B. R., & Rossion, B. (2015). The steady-state visual evoked potential in vision research: A review. Journal of Vision, 15(6), 4. https://doi.org/10.1167/15.6.4

Oostenveld, R., & Praamstra, P. (2001). The five percent electrode system for high-resolution EEG and ERP measurements. Clinical Neurophysiology, 112(4), 713–719. https://doi.org/10.1016/S1388-2457(00)00527-7

Ossandon, J. P., Helo, A. V., Montefusco-Siegmund, R., & Maldonado, P. E. (2010). Superposition Model Predicts EEG Occipital Activity during Free Viewing of Natural Scenes. Journal of Neuroscience, 30(13), 4787–4795.https://doi.org/10.1523/JNEUROSCI.5769-09.2010

Otero-Millan, J., Troncoso, X. G., Macknik, S. L., Serrano-Pedraza, I., & Martinez-Conde, S. (2008). Saccades and microsaccades during visual fixation, exploration, and search: Foundations for a common saccadic generator. Journal of Vision, 8(14), 21–21.https://doi.org/10.1167/8.14.21

Palomba, D., Angrilli, A., & Mini, A. (1997). Visual evoked potentials, heart rate responses and memory to emotional pictorial stimuli. International Journal of Psychophysiology, 27(1), 55–67. https://doi.org/10.1016/S0167-8760(97)00751-4

Patrick, C. J., Bradley, M. M., & Lang, P. J. (1993). Emotion in the criminal psychopath: Startle reflex modulation. Journal of Abnormal Psychology, 102(1), 82–92.https://doi.org/10.1037/0021-843X.102.1.82

Pedroni, A., Bahreini, A., & Langer, N. (2019). Automagic: Standardized preprocessing of big EEG data. NeuroImage, 200, 460–473. https://doi.org/10.1016/j.neuroimage.2019.06.046

Peirce, J. W. (2007). PsychoPy—Psychophysics software in Python. Journal of Neuroscience Methods, 162(1-2), 8–13. https://doi.org/10.1016/j.jneumeth.2006.11.017

Pernet, C., Garrido, M. I., Gramfort, A., Maurits, N., Michel, C. M., Pang, E., Salmelin, R., Schoffelen, J. M., Valdes-Sosa, P. A., & Puce, A. (2020). Issues and recommendations from the OHBM COBIDAS MEEG committee for reproducible EEG and MEG research. Nature Neuroscience. https://doi.org/10.1038/s41593-020-00709-0

Pernet, C. R., Appelhoff, S., Gorgolewski, K. J., Flandin, G., Phillips, C., Delorme, A., & Oostenveld, R. (2019). EEG-BIDS, an extension to the brain imaging data structure for electroencephalography. Scientific Data, 6(1). https://doi.org/10.1038/s41597-019-0104-8

Pernet, C. R., Garrido, M., Gramfort, A., Maurits, N., Michel, C., Pang, E., Salmelin, R., Schoffelen, J. M., Valdes-Sosa, P. A., & Puce, A. (2018). Best Practices in Data Analysis and Sharing in Neuroimaging using MEEG (preprint). Open Science Framework. https://doi.org/10.31219/osf.io/a8dhx

Picton, T. W. (2011). Human auditory evoked potentials [OCLC: 1097163932]. Plural Publishing Inc.

Picton, T. W., John, M. S., Dimitrijevic, A., & Purcell, D. (2003). Human auditory steady-state responses: Respuestas auditivas de estado estable en humanos. International Journal of Audiology, 42(4), 177–219.https://doi.org/10.3109/14992020309101316

Plöchl, M., Ossandón, J. P., & König, P. (2012). Combining EEG and eye tracking: Identification, characterization, and correction of eye movement artifacts in electroencephalographic data. Frontiers in Human Neuroscience, 6. https://doi.org/10.3389/fnhum.2012.00278

Poulsen, A. T., Kamronn, S., Dmochowski, J., Parra, L. C., & Hansen, L. K. (2017). EEG in the classroom: Synchronised neural recordings during video presentation. Scientific Reports, 7(1). https://doi.org/10.1038/srep43916

Quax, S. C., Dijkstra, N., van Staveren, M. J., Bosch, S. E., & van Gerven, M. A. (2019). Eye movements explain decodability during perception and cued attention in MEG. NeuroImage, 195, 444–453. https://doi.org/10.1016/j.neuroimage.2019.03.069

Raffaelli, Q., Mills, C., & Christoff, K. (2018). The knowns and unknowns of boredom: A review of the literature. Experimental Brain Research, 236(9), 2451–2462.https://doi.org/10.1007/s00221-017-4922-7

Rammstedt, B., Danner, D., Soto, C. J., & John, O. P. (2020). Validation of the Short and Extra-Short Forms of the Big Five Inventory-2 (BFI-2) and Their German Adaptations. European Journal of Psychological Assessment, 36(1), 149–161.https://doi.org/10.1027/1015-5759/a000481

Ramos Gameiro, R., Kaspar, K., König, S. U., Nordholt, S., & König, P. (2017). Exploration and Exploitation in Natural Viewing Behavior. Scientific Reports, 7(1), 2311. https://doi.org/10.1038/s41598-017-02526-1

Robbins, K. A., Touryan, J., Mullen, T., Kothe, C., & Bigdely-Shamlo, N. (2020). How Sensitive Are EEG Results to Preprocessing Methods: A Benchmarking Study. IEEE Transactions on Neural Systems and Rehabilitation Engineering, 28(5), 1081–1090. https://doi.org/10.1109/TNSRE.2020.2980223

Rosenthal, R. (1966). Experimenter effects in behavioral research. Appleton-Century-Crofts.

Roth, H. L. (2002). Effects of monocular viewing and eye dominance on spatial attention. Brain, 125(9), 2023–2035.https://doi.org/10.1093/brain/awf210

Saupe, K., Widmann, A., Bendixen, A., Müller, M. M., & Schröger, E. (2009). Effects of intermodal attention on the auditory steady-state response and the event-related potential. Psychophysiology, 46(2), 321–327. https://doi.org/10.1111/j.1469-8986.2008.00765.x

Schandry, R., Sparrer, B., & Weitkunat, R. (1986). From the heart to the brain: A study of heartbeat contingent scalp potentials. International Journal of Neuroscience, 30(4), 261–275.https://doi.org/10.3109/00207458608985677

Seabold, S., & Perktold, J. (2010). Statsmodels: Econometric and statistical modeling with python. 9th Python in Science Conference.

Silvia, P. J. (2009). Looking past pleasure: Anger, confusion, disgust, pride, surprise, and other unusual aesthetic emotions. Psychology of Aesthetics, Creativity, and the Arts, 3(1), 48–51.https://doi.org/10.1037/a0014632

Skosnik, P. D., Krishnan, G. P., & O’Donnell, B. F. (2007). The effect of selective attention on the gamma-band auditory steady-state response. Neuroscience Letters, 420(3), 223–228. https://doi.org/10.1016/j.neulet.2007.04.072

Smallwood, J., Beach, E., Schooler, J. W., & Handy, T. C. (2008). Going AWOL in the Brain: Mind Wandering Reduces Cortical Analysis of External Events. Journal of Cognitive Neuroscience, 20(3), 458–469.https://doi.org/10.1162/jocn.2008.20037

Smallwood, J., Davies, J. B., Heim, D., Finnigan, F., Sudberry, M., O’Connor, R., & Obonsawin, M. (2004). Subjective experience and the attentional lapse: Task engagement and disengagement during sustained attention. Consciousness and Cognition, 13(4), 657–690.https://doi.org/10.1016/j.concog.2004.06.003

Smith, T., & Henderson, J. (2010). Attentional synchrony in static and dynamic scenes. Journal of Vision, 8(6), 773–773.https://doi.org/10.1167/8.6.773

Smith, T. J., & Mital, P. K. (2013). Attentional synchrony and the influence of viewing task on gaze behavior in static and dynamic scenes. Journal of Vision, 13(8), 16–16.https://doi.org/10.1167/13.8.16

Smith, T. J. (2013). Watching You Watch Movies: Using Eye Tracking to Inform Cognitive Film Theory. In A. P. Shimamura (Ed.), Psychocinematics: Exploring cognition at the movies (p. 54). Oxford Univ. Press.

Snaith, R. P., Hamilton, M., Morley, S., Humayan, A., Hargreaves, D., & Trigwell, P. (1995). A scale for the assessment of hedonic tone the Snaith–Hamilton Pleasure Scale. British Journal of Psychiatry, 167(1), 99–103.https://doi.org/10.1192/bjp.167.1.99

Soto, C. J., & John, O. P. (2017). Short and extra-short forms of the Big Five Inventory–2: The BFI-2-S and BFI-2-XS. Journal of Research in Personality, 68, 69–81. https://doi.org/10.1016/j.jrp.2017.02.004

Stancin, I., Cifrek, M., & Jovic, A. (2021). A review of EEG signal features and their application in driver drowsiness detection systems. Sensors, 21(11), 3786. https://doi.org/10.3390/s21113786

Stapells, D. R., Linden, D., Suffield, J. B., Hamel, G., & Picton, T. W. (1984). Human Auditory Steady State Potentials: Ear and Hearing, 5(2), 105–113.https://doi.org/10.1097/00003446-198403000-00009

Stern, J. A., Boyer, D., & Schroeder, D. (1994). Blink Rate: A Possible Measure of Fatigue. Human Factors: The Journal of the Human Factors and Ergonomics Society, 36(2), 285–297.https://doi.org/10.1177/001872089403600209

Struk, A. A., Carriere, J. S. A., Cheyne, J. A., & Danckert, J. (2017). A Short Boredom Proneness Scale: Development and Psychometric Properties. Assessment, 24(3), 346–359.https://doi.org/10.1177/1073191115609996

Team, R. C. (2018). R: A language and environment for statistical computing.

Thielen, J., Bosch, S. E., van Leeuwen, T. M., van Gerven, M. A. J., & van Lier, R. (2019). Evidence for confounding eye movements under attempted fixation and active viewing in cognitive neuroscience. Scientific Reports, 9(1). https://doi.org/10.1038/s41598-019-54018-z

Thompson, E. R. (2007). Development and Validation of an Internationally Reliable Short-Form of the Positive and Negative Affect Schedule (PANAS). Journal of Cross-Cultural Psychology, 38(2), 227–242.https://doi.org/10.1177/0022022106297301

Tinio, P. P. L., Smith, J. K., & Smith, L. F. (2013). The walls do speak: Psychological aesthetics and the museum experience. In P. P. L. Tinio & J. K. Smith (Eds.), The Cambridge Handbook of the Psychology of Aesthetics and the Arts (pp. 195–218). Cambridge University Press. https://doi.org/10.1017/CBO9781139207058.011

Vallat, R. (2018). Pingouin: Statistics in Python. Journal of Open Source Software, 3(31), 1026. https://doi.org/10.21105/joss.01026

Van Eeckhoutte, M., Luke, R., Wouters, J., & Francart, T. (2018). Stability of Auditory Steady State Responses Over Time: Ear and Hearing, 39(2), 260–268. https://doi.org/10.1097/AUD.0000000000000483

van Gent, P., Farah, H., van Nes, N., & van Arem, B. (2019). HeartPy: A novel heart rate algorithm for the analysis of noisy signals. Transportation Research Part F: Traffic Psychology and Behaviour, 66, 368–378. https://doi.org/10.1016/j.trf.2019.09.015

Vessel, E. A. (2020). Neuroaesthetics. Reference Module in Neuroscience and Biobehavioral Psychology. Elsevier. https://doi.org/10.1016/B978-0-12-809324-5.24104-7

Vessel, E. A., Isik, A. I., Belfi, A. M., Stahl, J. L., & Starr, G. G. (2019). The default-mode network represents aesthetic appeal that generalizes across visual domains. Proceedings of the National Academy of Sciences, 116(38), 19155–19164.https://doi.org/10.1073/pnas.1902650116

Vessel, E. A., Maurer, N., Denker, A. H., & Starr, G. G. (2018). Stronger shared taste for natural aesthetic domains than for artifacts of human culture. Cognition, 179, 121–131. https://doi.org/10.1016/j.cognition.2018.06.009

Vessel, E. A., & Rubin, N. (2010). Beauty and the beholder: Highly individual taste for abstract, but not real-world images. Journal of Vision, 10(2), 1–14.https://doi.org/10.1167/10.2.18

Vessel, E. A., Starr, G. G., & Rubin, N. (2012). The brain on art: Intense aesthetic experience activates the default mode network. Frontiers in Human Neuroscience, 6. https://doi.org/10.3389/fnhum.2012.00066

Vodrahalli, K., Chen, P.-H., Liang, Y., Baldassano, C., Chen, J., Yong, E., Honey, C., Hasson, U., Ramadge, P., Norman, K. A., & Arora, S. (2018). Mapping between fMRI responses to movies and their natural language annotations. NeuroImage, 180, 223–231. https://doi.org/10.1016/j.neuroimage.2017.06.042

Vrana, S. R., Spence, E. L., & Lang, P. J. (1988). The startle probe response: A new measure of emotion? Journal of Abnormal Psychology, 97(4), 487–491.https://doi.org/10.1037/0021-843X.97.4.487

Waskom, M. (2021). Seaborn: Statistical data visualization. Journal of Open Source Software, 6(60), 3021. https://doi.org/10.21105/joss.03021

Winkler, I., Haufe, S., & Tangermann, M. (2011). Automatic Classification of Artifactual ICA-Components for Artifact Removal in EEG Signals. Behavioral and Brain Functions, 7(1), 30. https://doi.org/10.1186/1744-9081-7-30

Winton, W. M., Putnam, L. E., & Krauss, R. M. (1984). Facial and autonomic manifestations of the dimensional structure of emotion. Journal of Experimental Social Psychology, 20(3), 195–216. https://doi.org/10.1016/0022-1031(84)90047-7

Wong, P. K., & Bickford, R. G. (1980). Brain stem auditory evoked potentials: The use of noise estimate. Electroencephalography and Clinical Neurophysiology, 50(1-2), 25–34. https://doi.org/10.1016/0013-4694(80)90320-X

Yuval-Greenberg, S., Tomer, O., Keren, A. S., Nelken, I., & Deouell, L. Y. (2008). Transient Induced Gamma-Band Response in EEG as a Manifestation of Miniature Saccades. Neuron, 58(3), 429–441.https://doi.org/10.1016/j.neuron.2008.03.027

